# Arg-tRNA synthetase links inflammatory metabolism to RNA splicing and nuclear trafficking via SRRM2

**DOI:** 10.1101/2021.09.07.459304

**Authors:** Haissi Cui, Jolene K. Diedrich, Douglas C. Wu, Justin J. Lim, Ryan M. Nottingham, James J. Moresco, John R. Yates, Benjamin J. Blencowe, Alan M. Lambowitz, Paul Schimmel

## Abstract

Cells respond to perturbations like inflammation by sensing changes in metabolite levels. Especially prominent is arginine, which has known connections to the inflammatory response. Here, we found that depletion of arginine during inflammation decreased levels of a nuclear form of arginyl-tRNA synthetase (ArgRS). Surprisingly, we found that nuclear ArgRS interacts with serine/arginine repetitive matrix protein 2 (SRRM2), a spliceosomal protein and nuclear speckle component and that arginine depletion impacted both condensate-like nuclear trafficking of SRRM2 and splice-site usage in certain genes. These splice-site usage changes cumulated in synthesis of different protein isoforms that altered cellular metabolism and peptide presentation to immune cells. Our findings uncover a novel mechanism whereby a tRNA synthetase cognate to a key amino acid that is metabolically controlled during inflammation modulates the splicing machinery.

## Introduction

Inflammation accompanies most diseases, including cancer, neurodegeneration and auto-immune disorders (Netea et al., 2017). One of the metabolic hallmarks of inflammation is a decrease in systemic arginine levels due to its increased consumption by pro- and anti-inflammatory enzymes, such as iNOS and Arginase-1, with plasma arginine recovering when the inflammatory insult is resolved (Bronte and Zanovello, 2005; Murray, 2016). Consequently, arginine and signaling molecules derived from arginine *(e.g*., nitric oxide and ornithine) are key regulators of cellular signaling pathways, including the central metabolic regulator mTOR, which is necessary for immune cell activation during inflammation (Bar-Peled and Sabatini, 2014; Weichhart et al., 2015; Jones and Pearce, 2017). In addition to its physiological decrease during inflammation, therapeutic arginine depletion by recombinant enzymes is in clinical trials for cancer treatment (Patil et al., 2016).

The function of aminoacyl-tRNA synthetases (aaRSs) in cell signal transduction has come into focus in recent years. tRNA synthetases are highly conserved proteins that are essential for mRNA translation, where they catalyze the covalent attachment of amino acids to the 3’ end of their cognate tRNAs (Schimmel and Söll, 1979). Eight aaRSs with nine distinct catalytic activities together with three adapter proteins form the multisynthetase complex (MSC). While the MSC does not contribute to protein synthesis, complex formation is crucial for the intracellular distribution of certain aaRSs and specifically for their nuclear localization (Cui et al., 2021). The MSC is dynamic and acts as a reservoir to release aaRSs for extra-translational functions (Lee et al., 2004; Ray et al., 2007; Duchon et al., 2017).

aaRSs play a prominent role in mediating extracellular and nuclear cell signaling pathways. Although nuclear-localized aaRSs were historically seen as quality control agents for the correct processing of nascent tRNAs before their export to the cytoplasm (Lund and Dahlberg, 1998; Sarkar et al., 1999), they have also been linked to other functions (Guo and Schimmel, 2013; Shi et al., 2017; Kwon et al., 2019), including mediation of small molecule sensing (Sajish and Schimmel, 2015) and regulation of gene expression (Yannay-Cohen et al., 2009; Shi et al., 2014). In addition to freely diffusing nuclear aaRSs, the MSC is also found in the nucleus (Kaminska et al., 2009; Nathanson and Deutscher, 2000).

The cell nucleus is compartmentalized by membraneless organelles, some of which display condensate-like behavior, with the characteristics of phase-separated liquids (Shin and Brangwynne, 2017; Hnisz et al., 2017). Despite their lack of a membrane and constant exchange with the surrounding nucleoplasm, these subnuclear domains display structural integrity (Zhu and Brangwynne, 2015). In the nucleus, DNA-sparse interchromatin granule clusters (termed nuclear speckles) sequester proteins with repetitive sequences of low complexity and high disorder, among them factors necessary for mRNA maturation and mRNA splicing (Galganski et al., 2017; Saitoh et al., 2004; Spector and Lamond, 2011). RNA polymerases and splicing factors are in steady exchange between these structures and the surrounding nucleoplasm (Guo et al., 2019), raising the possibility that nuclear protein trafficking regulates their functions.

Different protein isoforms can be derived from the same coding sequence by alternative splicing, potentially resulting in different functions (Galarza-Muñoz et al., 2017). Alternative splicing events leading to intron retention can post-transcriptionally regulate mRNA levels by impeding transport to the cytoplasm and increasing RNA degradation (Braunschweig et al., 2014). 20% of alternatively spliced cassette exons (where splice sites are variably recognized leading to an exon being either retained or spliced out) are evolutionarily conserved between human and mouse, suggesting functional consequences (Pan et al., 2004).

Nuclear speckles have been suggested to store mRNA processing factors at their periphery and/or be hubs for mRNA transcription and processing (Chen and Belmont, 2019; Smith et al., 2020). A series of studies identified SRRM2 (serine/arginine repetitive matrix protein 2, also known as SRm300, gene name *SRRM2*), a component of spliceosomal complexes (Blencowe et al., 1998, 2000; Zhang et al., 2017), and SON, a mRNA binding protein and splicing factor, as being required for nuclear speckle formation (Miyagawa et al., 2012; Hu et al., 2019; Ilik et al., 2020). During mRNA splicing, the conserved N-terminal domain of SRRM2 engages with exon sequences and regulates splice-site selection (Gautam et al., 2015; Zhang et al., 2017). In line with its importance, SRRM2 mutations have been identified in developmental disorders (Kaplanis et al., 2020), cancer (Tomsic et al., 2015) and Parkinson’s disease (Shehadeh et al., 2010).

Given the role of arginine metabolism in inflammation (Bronte and Zanovello, 2005), we initiated our studies of the physiological role of nuclear aaRSs with arginyl-tRNA synthetase (ArgRS, gene name: *RARS*). We delineated a pathway through which cells respond to arginine starvation during inflammation via an interaction between ArgRS and SRRM2 that impacts trafficking of SRRM2 between nuclear speckles and the nucleoplasm and results in changes in alternative mRNA splicing. These alternative splicing changes culminated in altered cellular metabolism and peptide presentation to the immune system. Our results demonstrate a novel mechanism in which an aaRS interacts with a splicing factor to regulate nuclear trafficking and alternative mRNA splicing in response to a metabolic signal.

## Results

### Arginine promotes localization of arginyl-tRNA synthetase to the nuclear fraction

Previous work showed that ArgRS localizes to the nucleus as part of the MSC (Nathanson and Deutscher, 2000). To begin, we re-investigated the nuclear localization of ArgRS using four model systems: (i) human hepatic carcinoma cell line HepG2, (ii) human embryonic kidney endothelial cell line 293T, (iii) murine embryonic fibroblasts, and (iv) mouse liver. We confirmed the presence of ArgRS in the nuclear fraction of all four samples (Figure 1A, Supplementary Figure S1A). As arginine levels regulate central cellular signaling pathways (Bronte and Zanovello, 2005; Saxton et al., 2016), we tested whether nuclear ArgRS levels were affected by a reduction of extracellular arginine. We starved 293T cells of arginine for 6 h by incubation in arginine-free medium and found a decrease in the level of nuclear ArgRS, which could be reversed by adding arginine, while cytoplasmic ArgRS levels were unchanged by these treatments (Figure 1B).

**Figure 1:**
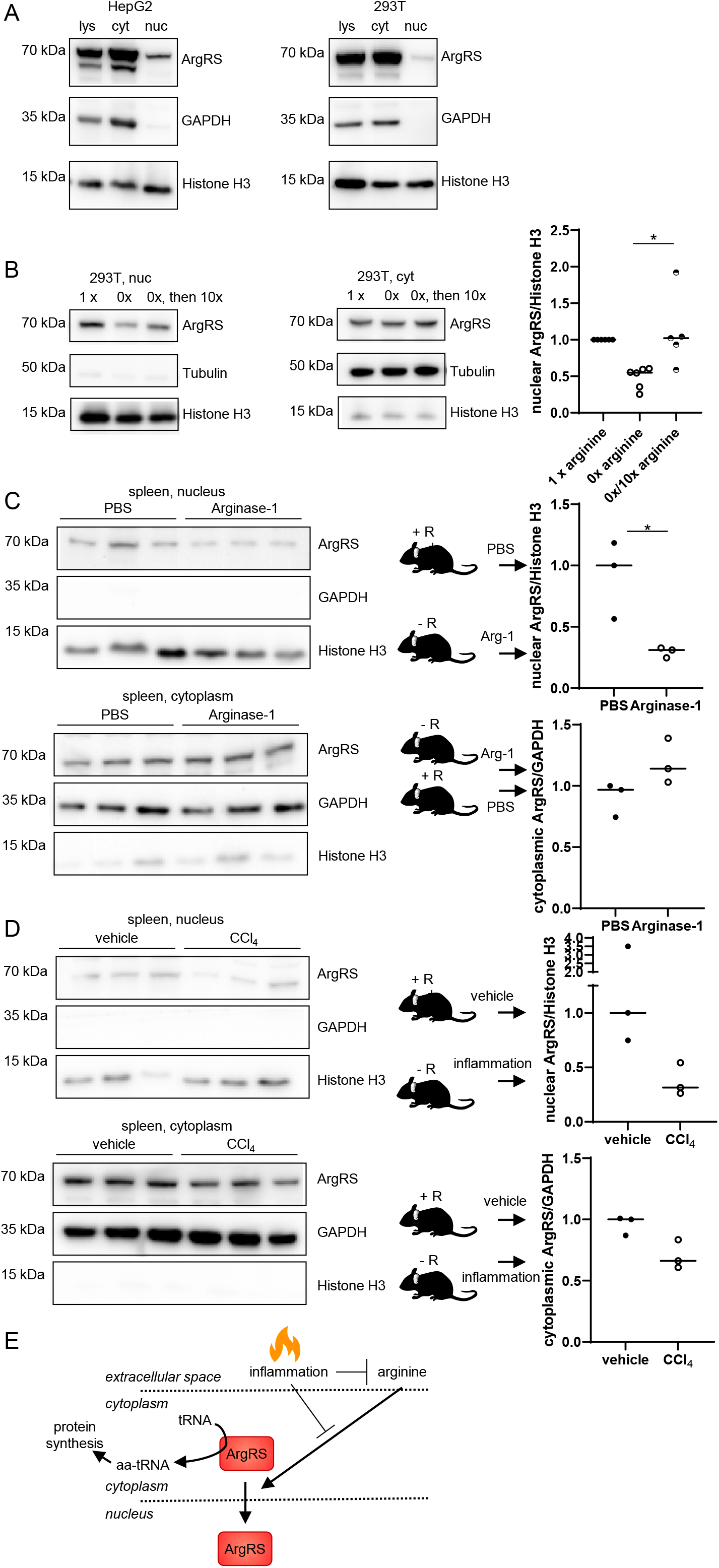
Arginyl-tRNA synthetase (ArgRS) localized to the nucleus in response to external arginine levels. (A) ArgRS in the cytoplasmic and nuclear fractions of human hepatocellular carcinoma HepG2 or HEK 293T cells. (B) ArgRS in the nucleus (left) and cytoplasm (right) following cell fractionation. Nuclear ArgRS could be restored by addition of external arginine after starvation (0x/10x). Arginine was re-added at 10x to DMEM to replenish lysosomal arginine storage and ensure sufficient arginine to rescue ArgRS nuclear localization. Arginine concentrations are relative to DMEM. p = *0.02. (C) Injection of recombinant Arginase-1 reduced nuclear ArgRS in the spleen of mice. p = *0.03. (D) Systemic inflammation by CCl_4_ reduced nuclear ArgRS in the spleen of mice. (A-D) Western blots and densitometric quantification. GAPDH: cytoplasmic marker, Tubulin: cytoplasmic marker, Histone H3: nuclear marker. (E) Scheme: Reduction of arginine during inflammation leads to reduced nuclear ArgRS.

### Arginine depletion and inflammation reduce nuclear ArgRS *in vivo*

Next, we verified the effect of arginine starvation on nuclear ArgRS levels in a more physiological model by reducing systemic arginine in mice by intraperitoneal injection of recombinant murine Arginase-1. Plasma arginine in the arginine-depleted mice was reduced to 30 - 50% of PBS controls (Supplementary Figure S1B), while nuclear ArgRS in the spleen, which consist mostly of immune cells, was reduced to < 40% of PBS controls (Figure 1C, p = 0.03).

As plasma arginine concentration was known to decrease during inflammation (Murray, 2016), we tested whether systemic inflammation in mice would similarly affect nuclear ArgRS levels by using the well-established liver toxin carbon tetrachloride (CCl_4_) (Horiguchi et al., 2010). Plasma arginine levels dropped to 50% of vehicle controls (Supplementary Figure S1C) and nuclear ArgRS levels in the spleen were reduced by ~60% (Figure 1D). These findings indicated that a decrease in systemic arginine by itself or as a result of inflammation induced by liver injury decreased nuclear ArgRS levels and focused our attention on the function of nuclear ArgRS (Figure 1E).

### The ArgRS interactome contains nuclear proteins

To identify interaction partners of nuclear ArgRS, which could offer clues to its function, we enriched ArgRS from the nuclear fraction of HepG2 cells using polyclonal ArgRS antibodies and characterized its protein interactome by using mass spectrometry (Keilhauer et al., 2015). We used label-free quantification (LFQ) to compare intensities against an antibody isotype control and calculated p-values from three replicates (Supplementary Table S1-4). Proteins with a p-value < 0.05 were identified as significantly enriched (Figure 2A). All MSC proteins, except leucyl-tRNA synthetase (LeuRS, gene name *LARS*), were retrieved from the HepG2 nuclear fraction (Figure 2A, orange, Supplementary Table S1), suggesting that LeuRS may be absent or sub-stoichiometric in the nuclear MSC. Aminoacyl tRNA Synthetase Complex Interacting Multifunctional Protein 3 (AIMP3) was enriched but with a p-value > 0.05. Thus, by enriching only for ArgRS, we captured nearly all MSC interactions. These results suggest that the MSC is mostly intact in the nucleus of HepG2 cells but may be in a more dynamic form than in the cytoplasm. They also confirm we can identify ArgRS interaction partners, even if the interactions are indirect and mediated through other proteins.

**Figure 2:**
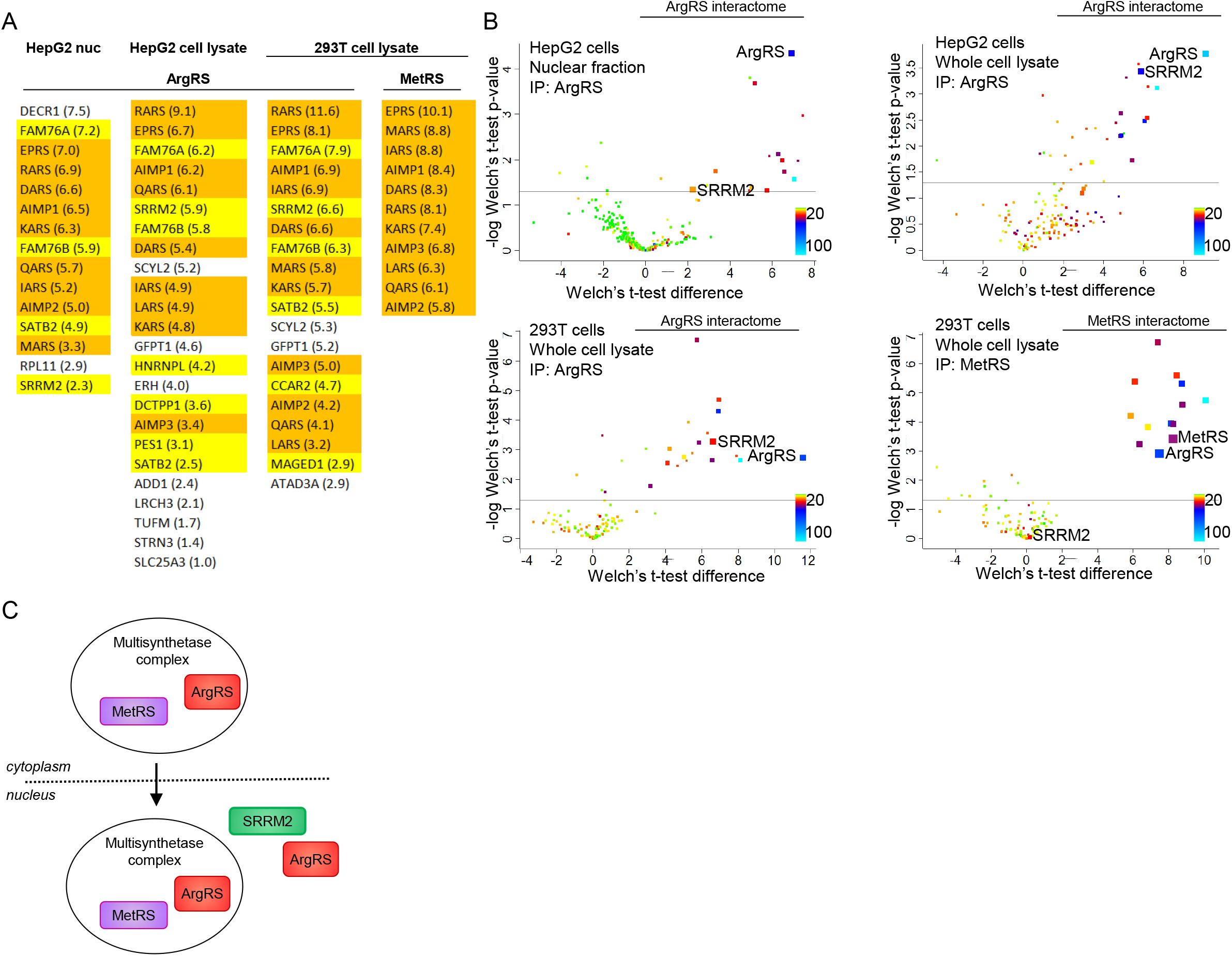
Characterization of the ArgRS interactome revealed nuclear proteins as interaction partners. (A) Comparison between the ArgRS interactome in nucleus, HepG2 whole cell lysate, 293T whole cell lysate, and with the interactome of MetRS, another tRNA synthetase. Multisynthetase complex (MSC) components: orange. Nuclear proteins: yellow. Proteins are identified by gene names and listed in order of enrichment over a non-targeted antibody of the same isotype with fold-change noted in parentheses. (B) Volcano plots of ArgRS or MetRS interactomes. MSC proteins are highlighted by larger symbol size. Axes denote enrichment (Welch’s t-test enrichment, x-axis) and reproducibility between 3 repeats (Welch’s t-test significance, y-axis). Color: number of unique peptides detected. Line: p-value = 0.05 (C) Scheme: The MSC is intact in the nucleus and ArgRS interacts with SRRM2 independent of the MSC.

Besides MSC components, we found several proteins with predominantly nuclear localization in the nuclear ArgRS interactome, including the nuclear speckle and spliceosome component SRRM2 (Figure 2A, yellow). To avoid artifacts due to the isolation of nuclei, we repeated the experiment using whole cell lysates. Specific and reproducible binding (p-value < 0.05) to nuclear proteins were confirmed (Figure 2A, Supplementary Table S2), despite ArgRS being a predominantly cytoplasmic protein. Consistent with the known cytoplasmic composition of the MSC (Kaminska et al., 2009), LeuRS was retrieved by ArgRS immunoprecipitation from the whole cell lysates of HepG2 cells (Figure 2A).

To test if these interactions are limited to the liver carcinoma cell line HepG2, we used the same approach on 293T cells (Figure 2A, Supplementary Table S3). Affinity enrichment of ArgRS retrieved the MSC and several proteins with predominately nuclear localization (Figure 2A, B). Comparison between the ArgRS interactomes in the two cell types showed significant overlap but also distinct binding partners (Supplementary Figure S1D). In contrast, the same experimental procedure in 293T cell lysates with a polyclonal antibody specific to methionyl-tRNA synthetase (MetRS, gene name *MARS*), which is localized to another subcomplex within the MSC (Kim and Kim, 2020), yielded exclusively MSC proteins as significant interactors (p-value < 0.05, Figure 2A, Supplementary Table S4, Figure 2B, lower right). Among the ArgRS binding partners, SRRM2 was highly and reproducibly enriched in both 293T and HepG2 cells (Figure 2A, B, upper right and lower left, Supplementary Table S5).

### Reciprocal co-immunoprecipitation of ArgRS and SRRM2

We focused on SRRM2 because of its critical roles in both RNA splicing (Blencowe et al., 2000; Zhang et al., 2017; Bertram et al., 2020) and nuclear compartmentalization (Hu et al., 2019; Ilik et al., 2020; Miyagawa et al., 2012). We confirmed by reciprocal co-immunoprecipitation and western blotting in both HepG2 and 293T cell lysates that pulldown of ArgRS enriched for SRRM2 and vice versa (Supplementary Figure S1E, F). Similar enrichment was seen using different ArgRS and SRRM2 antibodies (Supplementary Figure S1G-I). By contrast, MetRS, another component of the nuclear MSC, was not enriched by SRRM2 pulldown (Supplementary Figure S1J). These findings, together with the lack of SRRM2 enrichment in the MetRS interactome (Figure 2B, lower right), indicated that ArgRS interacts with SRRM2 independently of the MSC (Figure 2C).

### ArgRS and SRRM2 localize in proximity to each other in the nucleus

To localize where ArgRS and SRRM2 might interact in intact cells, we used immunofluorescent staining of HepG2 cells followed by airyscan super-resolution microscopy. Staining of SRRM2 with a monoclonal antibody raised against a C-terminal peptide (Zanini et al., 2017) showed SRRM2 in chromatin-low nuclear structures (Figure 3A, Supplementary Figure S2A, E, F, S2F for staining specificity). 80% of SON, which together with SRRM2 is crucial for nuclear speckle formation (Ilik et al., 2020), colocalized with SRRM2 in chromatin-low areas (Supplementary Figure S2A), suggesting that these structures are indeed nuclear speckles.

**Figure 3:**
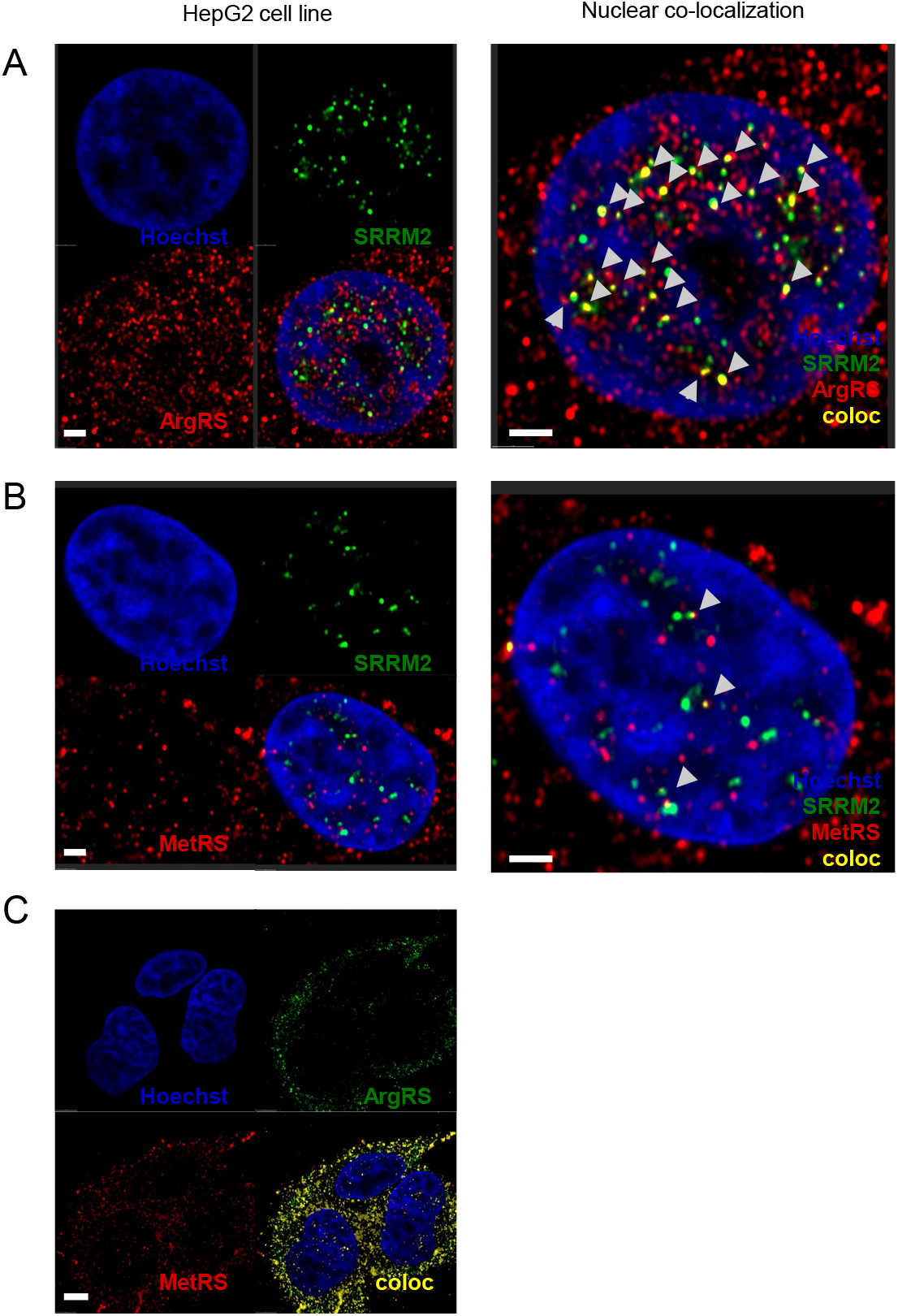
SRRM2 colocalized with ArgRS but not with MetRS. (A-C) Confocal microscopy following immunofluorescent staining of ArgRS/SRRM2 (A), MetRS/SRRM2 (B), or ArgRS/MetRS (C), respectively. A single slice of a series of z-stacks is shown. Colocalized areas in yellow. Bar: A, B: 2 μm. C: 5 μm.

We next assessed where ArgRS and SRRM2 localized. Immunofluorescence co-staining of ArgRS with a polyclonal antibody visualized ArgRS in and at the periphery of SRRM2-rich areas (Figure 3A, Supplementary Video S2, Supplementary Figure S2F for additional images, Supplementary Figure S2B for staining specificity). The calculated Manders overlap coefficient, which denotes the correlation of absolute intensities, was > 0.6 for SRRM2 protein within the threshold SRRM2 signal in ArgRS/SRRM2 co-staining (Supplementary Figure S2C). If the Costes method (Costes et al., 2004) was used for thresholding, colocalization increased to 0.8 (Supplementary Figure S2D). The Pearson correlation coefficient, another measure of colocalization which denotes the correlation of relative intensities, was 0.45 (Supplementary Figure S2I). About 60% of SRRM2 was calculated to be colocalized with ArgRS, which could be reproduced through co-staining with an alternative (monoclonal) ArgRS antibody together with a polyclonal SRRM2 antibody (Supplementary Figure S2E, Supplementary Figure S2F for additional images and staining specificity, Manders coefficient 0.6 (SRRM2)).

Nuclear ArgRS was also found outside of SRRM2-rich areas (Figure 2). We co-stained ArgRS with proteins located in other nuclear condensates structures to test whether ArgRS accumulated in other compartments or whether it was specific to SRRM2 and nuclear speckles. Co-staining of the paraspeckle protein SFPQ with ArgRS, yielded a Manders coefficient < 0.3 (Supplementary Figure S2G). As SFPQ was found throughout the nucleus, we also co-stained ArgRS with coilin, which exclusively located to Cajal bodies, and found a Manders coefficient close to 0 (Supplementary Figure S2H). Taken together, these results suggest ArgRS and SRRM2 specifically colocalize in the periphery of SRRM2-rich, low-chromatin regions identified as nuclear speckles. Arginine starvation significantly decreased the Pearson coefficient between ArgRS and SRRM2 (p = 0.01; Supplementary Figure S2I) from 0.6 to 0.4, in agreement with our finding that nuclear ArgRS levels were decreased by arginine starvation.

In accordance with our mass spectrometry results (Figure 2), the Manders coefficient for SRRM2/MetRS was only 0.13, as opposed to 0.65 for SRRM2/ArgRS (Figure 3B, Supplementary Figure 2C, Supplementary Video S3, p < 0.0001). If ArgRS interacts with SRRM2 while in the MSC, MetRS would be expected to colocalize with SRRM2 as well (Figure 3B). The majority of nuclear ArgRS did not co-stain with nuclear MetRS while cytoplasmic ArgRS and MetRS showed high colocalization (Manders coefficients 0.08 vs 0.72, Figure 3C, Supplementary Figure S2C, Supplementary Video S4), suggesting that the majority of nuclear ArgRS is not associated with the MSC. These results point to an ArgRS-specific, MSC-independent association with the nuclear speckle protein SRRM2 within or at the periphery of SRRM2-rich structures (Figure 3D).

### ArgRS depletion enhances SRRM2 mobility in nuclear speckles

ArgRS localized to the periphery of SRRM2-rich signals, so we wondered whether ArgRS might thereby influence SRRM2 mobility and trafficking. To address this question, we used fluorescence recovery after photobleaching (FRAP) to monitor SRRM2 flux (Figure 4A). For FRAP measurements, we fluorescently labeled endogenous SRRM2 by introducing an mVenus-tag four amino acids from the C-terminus of SRRM2 (Supplementary Figure S2J). As expected, SRRM2-mVenus in 293T cells localized to discrete nuclear regions similar to those detected by antibody-staining (Supplementary Figure S2K).

**Figure 4:**
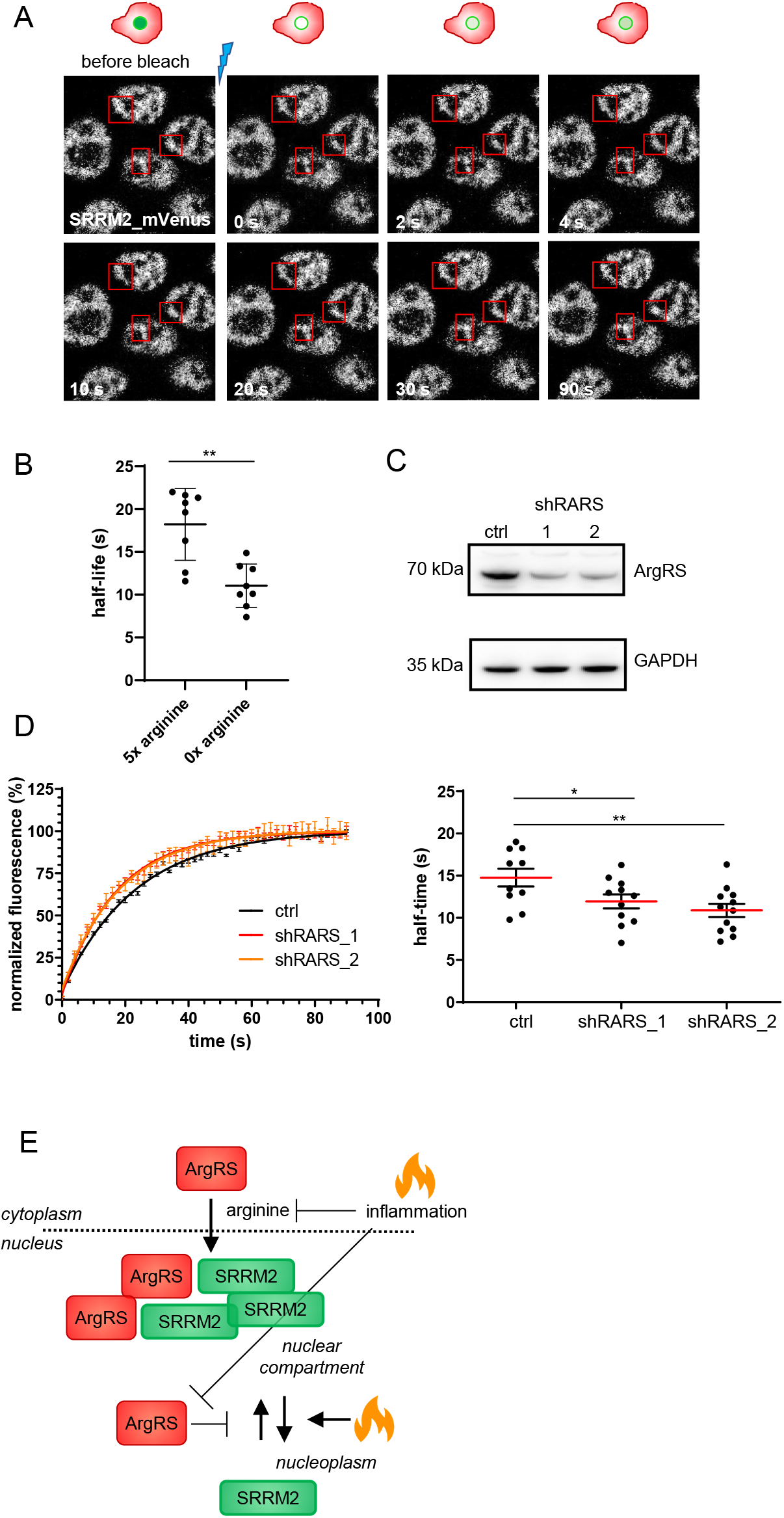
ArgRS knock down increased SRRM2 mobility. (A) Fluorescence recovery after photobleaching (FRAP). SRRM2-mVenus-dense areas were bleached, and recovery was monitored over 90 s. SRRM2-mVenus is displayed as greyscale. (B) Arginine starvation reduced SRRM2-mVenus FRAP half-time t_1/2_ to 60% as compared to high arginine. Arginine concentrations relative to DMEM, n = 8, p = **0.001. (C) Stable knock down of ArgRS in SRRM2-mVenus cell lines with two different shRNAs (shRARS_1, shRARS_2) reduced ArgRS protein levels to 40% of a non-targeting shRNA control, shown by western blot. Relative *RARS* mRNA levels quantified by qRT-PCR: shSCR: 1.00 ± 0.17; shRARS_1: 0.20 ± 0.17; shRARS_2: 0.07 ± 0.06. (D) Left panel: Recovery of fluorescent signal after photobleaching. Errors bars depict standard error of the mean. Right panel: t_1/2_ of SRRM2-mVenus recovery time after bleaching was reduced to 70% by stable ArgRS knock down. shSCR: n = 10, shRARS_1: n = 11, shRARS_2: n = 12, p = *0.048, **0.007. (E) Scheme: ArgRS impedes SRRM2 trafficking by slowing the exchange between nuclear compartments and nucleoplasm.

For FRAP experiments, regions of interest (ROIs) were chosen that contain one nuclear speckle including its surrounding nucleoplasm (Figure 4A). Bleaching of SRRM2-mVenus in ROIs was followed by a time-dependent recovery of fluorescence that plateaued within 60 s (Supplementary Figure S2L) through influx of SRRM2 from the surrounding SRRM2-rich compartments and nucleoplasm (Figure 4A, Supplementary Video S5-7). About 50% of SRRM2 in the ROIs was immobile (Supplementary Figure S2M). A one-exponential curve fit of background-subtracted, normalized fluorescent recordings estimated a recovery half-time of 17 s in arginine-rich medium (Figure 4B, D). SRRM2 therefore shows dynamic properties reminiscent of phase-separated molecules (Brangwynne et al., 2009).

This half-time of recovery is 5x longer than that for the nuclear speckle marker protein ASF/SRSF1 (3 s, Phair and Misteli, 2000). If SRRM2 and SRSF1 have dynamic movements that reflect simple diffusion kinetics, where the rate of diffusion varies inversely with the square root of the molecular weight, then the 10-fold higher molecular weight of SRRM2 would predict a 3x difference in their dynamics for spherical objects. From this perspective, given the disordered structure of both proteins, but especially of SRRM2, a 5-fold higher half-time of recovery seems reasonable. Arginine starvation, which decreased the amount of nuclear ArgRS (Figure 1B), increased SRRM2 mobility as indicated by the shorter half-time of recovery of SRRM2 (11 s vs 18 s) (Figure 4B, Supplementary Figure S2N).

The above findings suggested that nuclear ArgRS decreases SRRM2 mobility. To further investigate this possibility, we stably introduced shRNAs directed against the 3’ UTR (shRARS_1) or 5’ end (shRARS_2) of *RARS* mRNA. With both shRNAs, we reduced the ArgRS protein level in SRRM2-mVenus-expressing 293T cells to ~40% of that in cells expressing a non-targeting shRNA control (Figure 4C). This level of ArgRS knock down did not decrease cell viability or proliferation (Supplementary Figure S2O), in agreement with findings for knock downs of other aminoacyl-tRNA synthetases (Hayano et al., 2016; Ofir-Birin et al., 2013) and in a haplo-insufficient mouse (Seburn et al., 2006). SRRM2 protein expression levels were not reduced by the ArgRS knock downs (Supplementary Figure S2P). In both ArgRS knock down cell lines the half-time of recovery in SRRM2-mVenus regions decreased significantly to ~70% of the control (p = 0.048, 0.007) (Figure 4D, Supplementary Video S5-7), consistent with the increased mobility of SRRM2 seen for arginine starvation. Given that ArgRS is a 70 kDa protein and SRRM2 300 kDa (without the mVenus label), this decrease is in line with the expected effect size. Collectively, these findings indicate that ArgRS slowed SRRM2 speckle-trafficking, and that this interaction is modulated by the external arginine concentration, which in turn controls the amount of ArgRS in the nucleus (Figure 4E).

### ArgRS affects processing of mRNAs but not small ncRNAs

As SRRM2 is both a nuclear compartment organizer and spliceosome component, we hypothesized that arginine-mediated changes in the level of nuclear ArgRS, which impact SRRM2 mobility, could affect RNA processing. Because manipulation of arginine triggers several cross-talking cellular signaling pathways (Bar-Peled and Sabatini, 2014; Dong et al., 2000), we used ArgRS and SRRM2 knock downs as surrogates to assess how the interaction of SRRM2 with ArgRS might affect gene expression. For this purpose, we stably knocked down ArgRS in HepG2 cells (to ~ 50% on the protein level and ~ 40% on the RNA level, Supplementary Figure S3A, B) and reset the cellular response to arginine by 6 h of arginine starvation followed by 2 h of arginine supplementation. In previous experiments, we lowered total cellular ArgRS by 90% without affecting tRNA aminoacylation or mRNA translation (Cui et al., 2021). Consequently, the milder knock down of ArgRS to 50% was not expected to affect the housekeeping function of ArgRs in protein synthesis. We isolated RNA from a control shRNA-expressing cell line (shSCR ctrl) and one expressing a shRNA against ArgRS (shRARS) and confirmed that the knock down did not impair cell proliferation (Supplementary Figure S3C).

For a comprehensive view of the effect of ArgRS depletion on cellular RNA synthesis, we used TGIRT-seq of chemically fragmented, rRNA-depleted cellular RNAs. This method employs the novel template-switching activity of the TGIRT enzyme for 3′ RNA-seq adapter addition and enables profiling of coding and non-coding RNAs simultaneously (Figure 5A) (Qin et al., 2016; Nottingham et al., 2016; Boivin et al., 2018).

**Figure 5:**
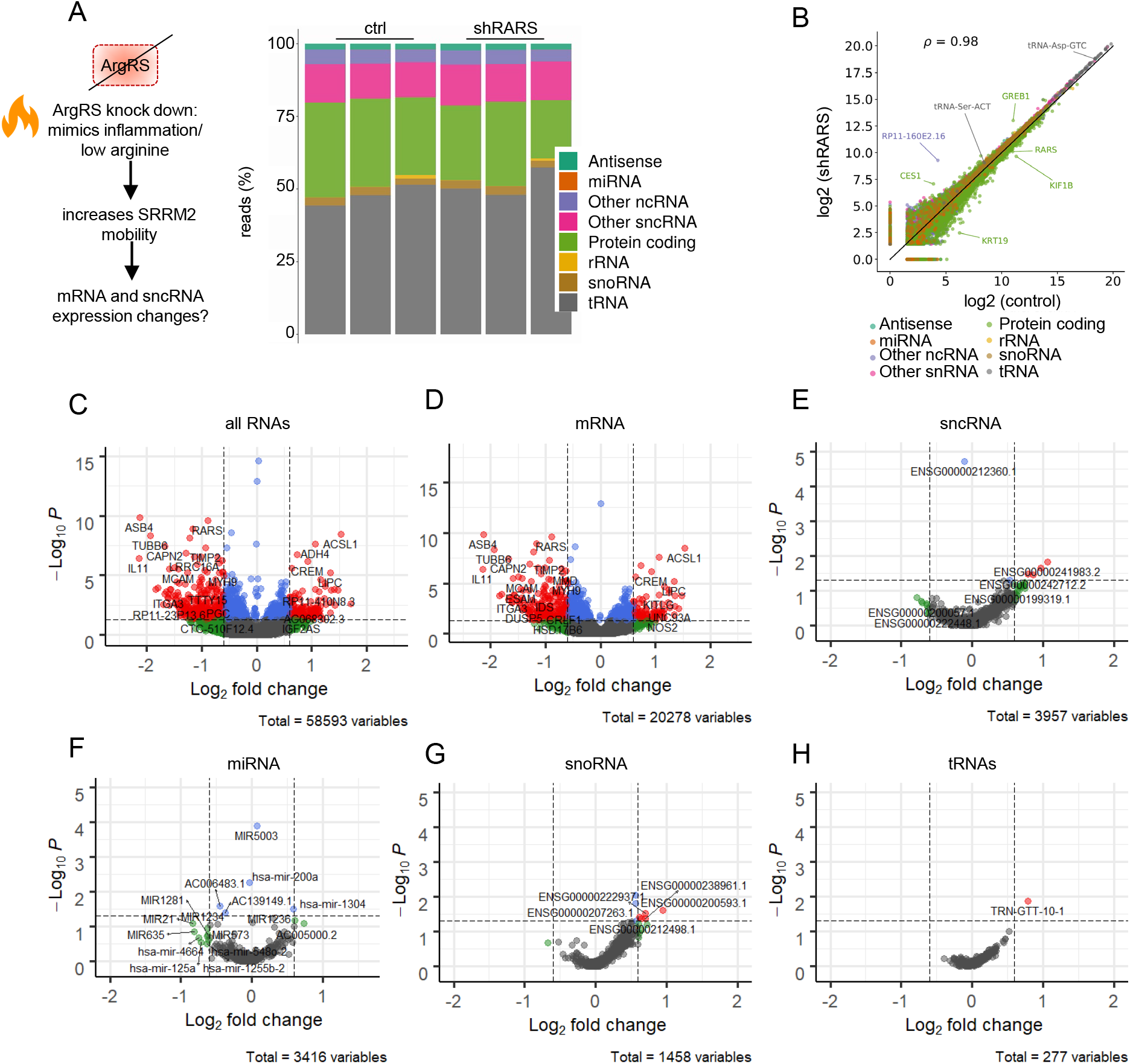
ArgRS knock down primarily affected mRNAs but not non-coding RNAs. (A) Stacked bar graphs showing RNA biotype distributions. (B) Scatter plot comparing RNAs. ρ is the Spearman’s correlation coefficient. (C-H) Volcano plots comparing all and different RNA biotypes in ArgRS knock down versus control HepG2 cells. Vertical dashed lines depict a 1.5-fold change and horizontal dashed line a padj < 0.05. Differential expression of (C) all RNAs, (D) mRNAs, (E) small non-coding RNAs, (F) microRNAs, (G) small nucleolar RNAs, and (H) transfer RNAs. (A-H): ctrl: HepG2 shSCR control. shRARS: HepG2 ArgRS knock down. Three replicate TGIRT-seq datasets.

Comparing 21,540 different coding and non-coding RNAs showed relatively little difference in the proportion of different RNA biotypes between the ArgRS shRNA knock down and control shRNA (Figure 5B). Most differentially expressed genes were protein-coding genes (Figure 5C, D). The ArgRS knock down also resulted in numerous differences in exon usage for these genes (Supplementary Figure S3D). In contrast, we found only minor differences in the levels of small ncRNAs (Figure 5E-H) with only a small change in the relative abundance of only one tRNA (tRNA^Asn^-GTT-10-1, p-value = 0.01) and no significant changes in the relative abundance of other tRNAs, including tRNA^Arg^ isodecoders (Figure 5H, Supplementary Table S6).

### ArgRS knock down alters mRNA processing and protein isoforms

Due to the higher representation of small ncRNAs, the sequencing depth we achieved with TGIRT-seq for mRNA exon-exon junctions was insufficient to draw conclusions about the significance of changes in splice junction usage. Thus, we prepared and sequenced libraries from the same RNA preparations after pre-selection for poly(A)-containing mRNAs. We used vast-tools (Tapial et al., 2017) to evaluate previously annotated splice junctions based on reads spanning exons (in the following referred to as “alternative splicing events”). We first identified alternative splicing events that were affected by ArgRS knock down compared to the shSCR control (Han et al., 2017; Irimia et al., 2014). For this purpose, we used the minimum value of the difference (denoted MV) as a measure of significance, where MV > 0 means a ≥ 0.95 probability that splice junction usage changed between the ArgRS knock down and the control (Han et al., 2017) (see Methods). These events were then further classified by |ΔPSI|, which is defined as the % difference spliced-in. We thus identified 393 events in 345 genes with alternative splicing changes between the ArgRS knock down and control with MV > 0 (Figure 6A shows events with |ΔPSI| > 0.1, Supplementary Table S7). These alternative splicing events corresponded predominantly to the inclusion or skipping of cassette exons and increased retained introns, while inclusion or skipping of microexons (3-27 nt) or use of alternative 5′ or 3′ splice sites were relatively unaffected by the ArgRS knock down (Fig. 6A).

**Figure 6:**
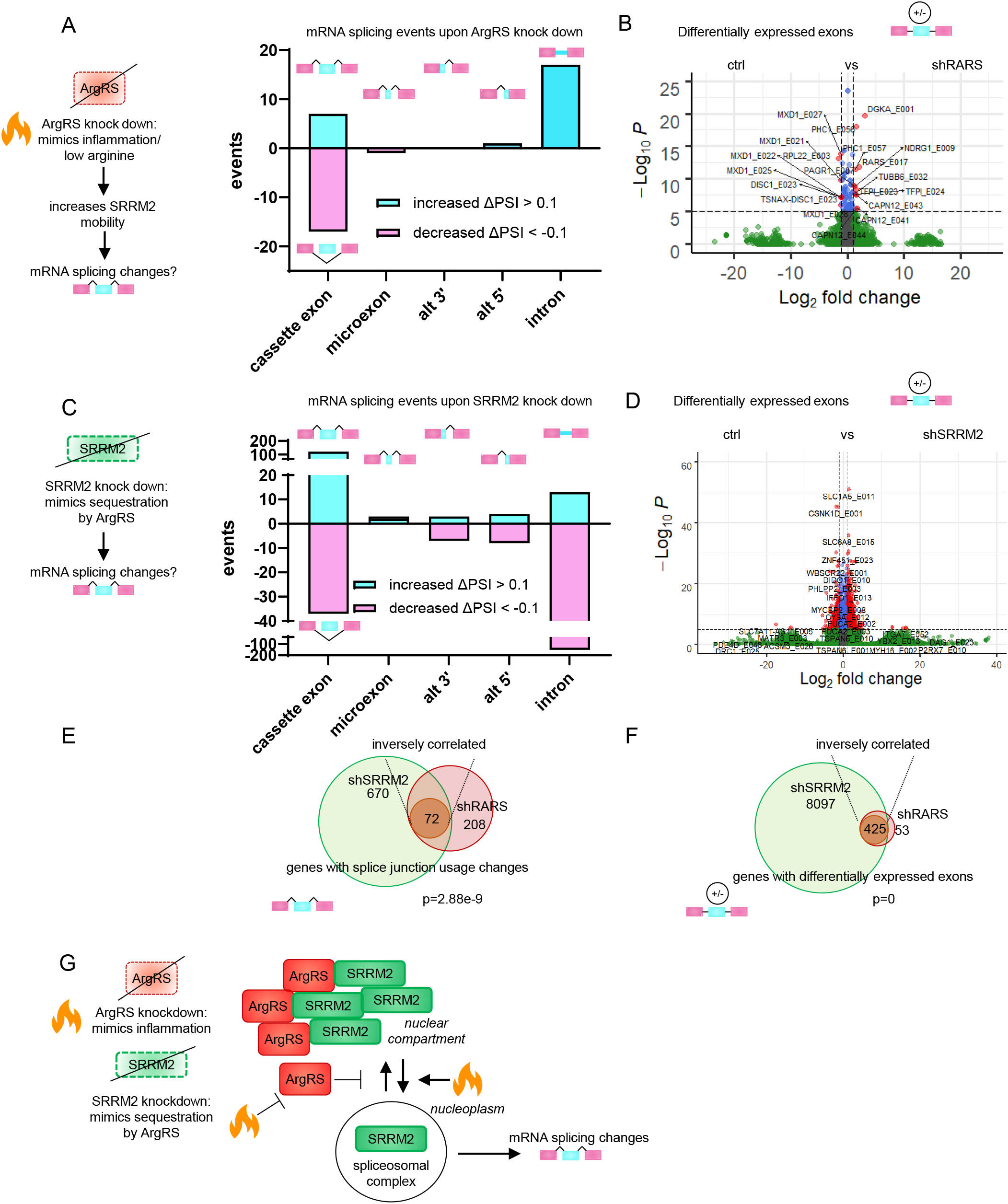
Differential mRNA processing upon ArgRS knock down. (A) Splicing event changes upon ArgRS knock down (shRARS). (B) Volcano plot of differentially expressed exons upon ArgRS knock down (shRARS). (C) Splicing event changes upon SRRM2 knock down. (D) Volcano plot of differential exon usage upon SRRM2 knock down. (A, C): Splicing events divided by event type, relative to shSCR control, as detected by RNA-seq. Events with a |ΔPSI| > 0.1, MV > 0 are shown. Increased event frequency shown in cyan, events with decreased frequency in pink. PSI: percent spliced in. MV > 0: significant splicing changes. Cassette: cassette exons/exon skipping, microexons: 3-15 nucleotide exons, alt 3’: alternative 3’ end splice site, alt 5’: alternative 5’ end splice site, intron: intron retention. 115814 (shRARS) and 111346 (shSRRM2) events were detected in total. (B, D) Dashed vertical lines: 2-fold change, dashed horizontal line: p-value 1e-6. (E, F) Venn diagram of genes with shared alternative splicing events or differentially expressed exons upon ArgRS and SRRM2 knock down. Genes with inversely regulated (E) splicing events (ΔPSI > 0 for shRARS and ΔPSI < 0 for shSRRM2 or vice versa) or (F) differentially expressed exons (log2fold change > 0 for shRARS or < 0 for shSRRM2 or vice versa) in brown circle. Hypergeometric test for enrichment of inversely correlated, ArgRS regulated splicing events. (A-F) Data was derived from sequencing of 3 replicates. All changes relative to shSCR control. (G) ArgRS slows down SRRM2 trafficking and thereby disrupts SRRM2-dependent mRNA splicing changes, leading to the regulation of alternative mRNA splicing in opposite directions by ArgRS and SRRM2.

We also analyzed differential transcript usage with DEXseq (Anders et al., 2012), which compares expression levels of individual exons to the rest of the gene as opposed to quantifying splice junction spanning reads. By using this complementary method, we identified 1,004 exons in 533 genes that were differentially expressed upon ArgRS knock down compared to the control (padj < 0.05, Figure 6B, Supplementary Table S8).

By using RT-PCR, we directly confirmed alternative splicing events upon ArgRS knock down for *FGFR3* and *PLEKHG5* in HepG2 cells (Supplementary Figure S4A, B). These genes displayed the same regulation in 293T cells upon ArgRS knock down (Supplementary Figure S4A, B). Rescue of ArgRS knock down by overexpression of a *RARS* transgene restored the ratio of *FGFR3* splice variants to that of control cells (Supplementary Figure S4C, D).

To investigate if the splicing changes upon ArgRS knockdown led to an altered proteome, we assessed splice variants using mass spectrometry. As proteins derived from alternative splicing can only be identified from individual peptides (Blencowe, 2017), we sought to increase the sensitivity of detection by enriching these peptides. Therefore, we focused on events where alternative splicing would give rise to distinct N-termini and enriched N-terminal peptides by removing all other (tryptic) peptides (Kleifeld et al., 2011). We identified 167 peptides from 135 proteins (Supplementary Table S9) that were detected at higher or lower intensity upon ArgRS knock down (|pdiff| > 1). Ten proteins were encoded by genes with differential exon usage and seven by genes with alternative splicing events upon ArgRS knockdown (Supplementary Figure S4E), including splicing changes that affected peptides in the proximity of potential alternative translation start sites (Supplementary Figure S4F). For the splicing factor PTBP1, we saw an increase of the N-terminal peptide for one isoform while the N-terminal peptide of another isoform was simultaneously reduced (Supplementary Figure S4F). Differences in PTBP1 isoform usage were confirmed by western blots (Supplementary Figure S4G). Collectively, these findings indicate that ArgRS knock down at levels that do not affect cell proliferation results in alternative splicing changes that lead to different protein isoforms.

### Identification of SRRM2-dependent splicing changes

Nuclear ArgRS interacts with and impedes SRRM2 mobility in the nucleus (Figure 4) and could thereby reduce the levels of SRRM2 available for RNA splicing. To identify alternative splicing events that are sensitive to decreasing levels of functional SRRM2, we knocked down SRRM2 by 50% (RNA and protein level, Supplementary Figure S5A). This knockdown led to 1,033 alternative splicing events in 807 genes with MV > 0 (Figure 6C shows events with |ΔPSI| > 0.1, Supplementary Table S10) and 42,140 differential exon usage events in 8,563 genes that were changed relative to the control shRNA (Figure 6D, Supplementary Table S11, padj < 0.05). The numbers of changed alternative splicing and differential exon usage events were much higher than those for the ArgRS knock down (393 and 1,004, respectively) (Figure 6A), as expected for a spliceosome component, whose activity might be subject to multiple modes of regulation. Cassette exons were changed in both directions upon SRRM2 knock down while introns were increasingly spliced out (Figure 6C). In order to rule out that the alternative splicing changes were due to off-target effects of the SRRM2 knock down with a single shRNA, we transfected 293T cells with a pool of siRNAs against SRRM2. Alternative splicing of *FGFR3* and *CDK5RAP3* genes could also be observed upon the reduction of SRRM2 with an heterogenous pool of siRNAs, suggesting that this was a direct consequence of lowered SRRM2 availability (Supplementary Figure S5B).

### ArgRS and SRRM2 modulate alternative splicing of a set of genes in opposite directions

After obtaining lists of genes that were sensitive to reduction of either SRRM2 or ArgRS, we next set out to identify which genes were shared between both. To control for indirect effects due to knockdown of an aaRSs, we also knocked down MetRS to 10% on RNA and protein level (which is lower than the ArgRS knock down, Figure S3A) as a control (Supplementary Figure S6A, Supplementary Table S12). We identified 602 alternative spicing events in 493 genes (|MV| > 0, events with |ΔPSI| > 0.1 shown in Supplementary Figure S6B) and 50,896 differentially expressed exons in 7,668 genes that changed upon MetRS knock down (Supplementary Figure S6B).

A total of 282 alternative splicing events overlapped between the ArgRS, SRRM2, and MetRS knock downs (Supplementary Figure S6C, Venn diagram, MV > 0, no criterion for |ΔPSI| to depict all events regardless of magnitude). Of these, 70 splicing events were shared between the SRRM2 and ArgRS knock downs (Supplementary Figure S6C) and their ΔPSIs were negatively correlated (R = −0.29, p = 0.037). We then compared the directionality of the overlapping splicing events between ArgRS and SRRM2 knock downs. This identified 27 of the 70 shared alternative splicing events (38.6%) that were inversely regulated by ArgRS and SRRM2, meaning that the knock down of ArgRS led to a ΔPSI > 0, while knock down of SRRM2 led to a ΔPSI < 0 or vice versa (Supplementary Figure S6D, inner brown circle, MV > 0, hypergeometric test for overrepresentation p = 0.01). In contrast, only 12/185 splicing events (6.2%) were inversely regulated by MetRS and SRRM2 (Supplementary Figure S6D, MV > 0, hypergeometric test p = 1). We confirmed splicing changes by RT-PCR for prominently changed events (*CDK5RAP3, IL32, RAD52*, and *STX16*, all Supplementary Figure S6E).

If focusing on genes that were alternatively spliced and shared between the ArgRS and SRRM2 knock downs (instead of individual splicing events within a gene that were shared), close to 500 genes had splicing events or differentially expressed exons that were inversely regulated by ArgRS and SRRM2 (72 and 475, respectively, Figure 6E, F). These findings suggested that the observed inverse regulation of alternative splicing by ArgRS and SRRM2 might lead to the production of different protein isoforms that contribute to the cellular response to inflammation (Figure 6G).

### ArgRS and SRRM2-regulated splicing alters cellular metabolism and communication

To explore the possible biological consequences of the ArgRS/SRRM2 interaction, we examined the enrichment of pathways and GO terms in the genes changed in opposite directions in the ArgRS and SRRM2 knock downs (Figure 7A, B, Supplementary Figure S7A-E). Among these were genes that might benefit the cellular response to inflammation such as the pro-inflammatory cytokine IL32 (Supplementary Figure S6E) (Aass et al., 2021). Overrepresented pathways included the mTORC1-pathway, which can be activated by arginine (Figure 7B).

**Figure 7:**
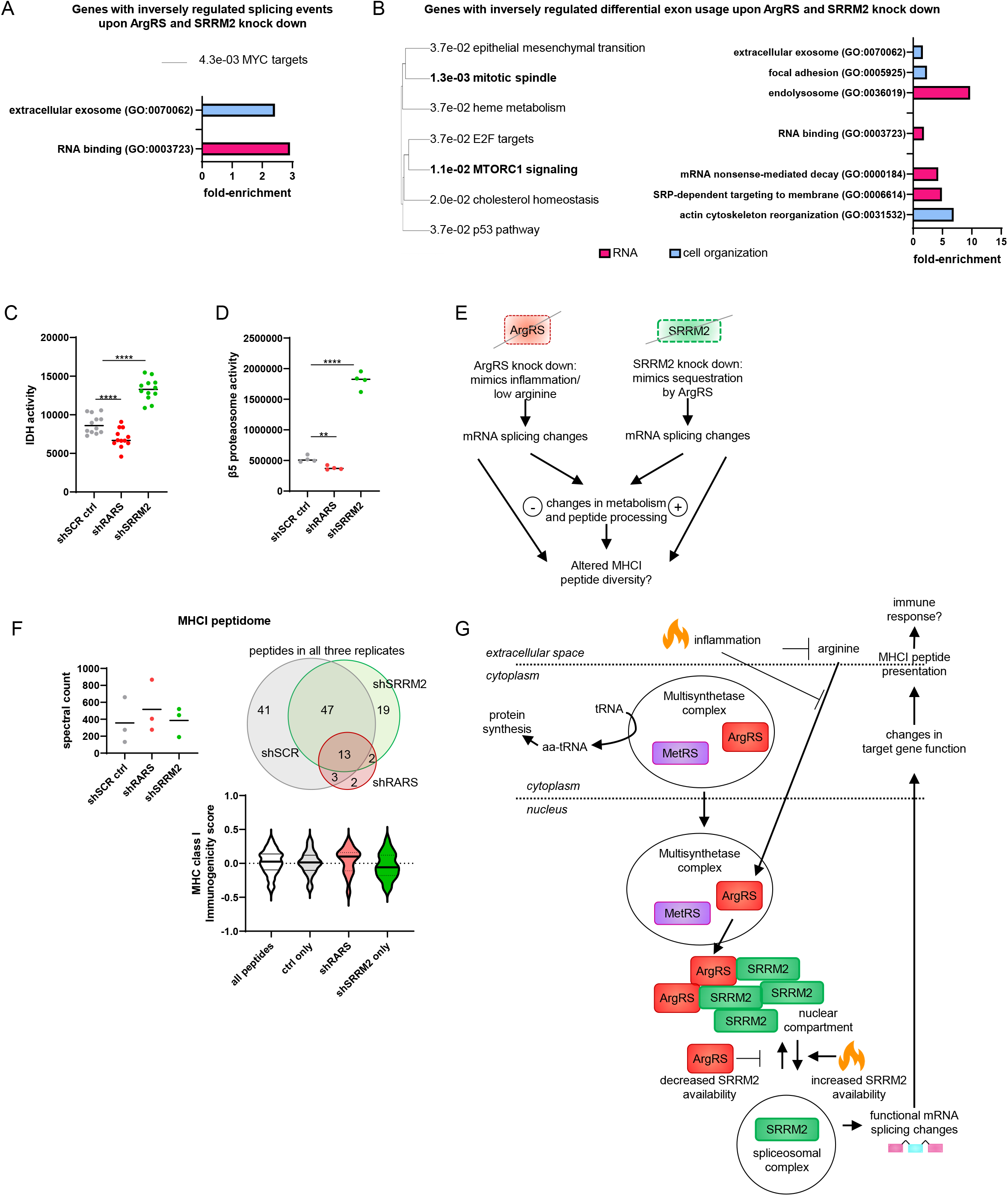
SRRM2- and ArgRS-regulated mRNA processing led to metabolic changes and an altered MHCI peptidome. (A) MSigDB and Gene Ontology (GO) analysis of genes with inversely regulated splicing events upon ArgRS and SRRM2 knock down. (B) MSigDB and GO analysis of genes with inversely regulated differentially expressed exons upon ArgRS and SRRM2 knock down. (A, B) GO terms associated with RNA-interacting proteins in fuchsia, GO terms associated with cellular organization and organelles in blue. (C) IDH1 enzyme activity. IDH activity assay: representative of 3 repeats, 12 technical replicates, p < ****0.0001. Deletion of ENSE00000934683 leads to an in-frame deletion of 126 amino acids. (D) PSMB5 differential exon usage, β5 proteasomal subunit (PSMB5 protein) activity. Quantification of differentially used exon: β5 activity: representative of 3 repeats, 4 replicates, p = **0.006, p < ****0.0001. (E) Model of ArgRS and SRRM2 knock down, their connection to inflammation, and consequences on their splice targets. (F) Total spectral counts (upper left) correspond to peptide quantity. Venn diagrams of identified MHCI peptides by mass spectrometry following isolation of MHCI complex (upper right) found in all replicates upon ArgRS or SRRM2 knock down. Bottom: Violin plot of MHCI immunogenicity score for peptides found in all samples, shSCR ctrl, ArgRS knock down, and SRRM2 knock down only. Median MHCI immunogenicity score is indicated by a black line. (A-F) shSCR ctrl: HepG2 shSCR control. shRARS: HepG2 ArgRS knock down. shSRRM2: HepG2 shSRRM2 knock down. (G) Model of cellular response to inflammation by arginine/ArgRS/SRRM2.

To investigate the functional consequences of differential exon usage that might be regulated by ArgRS/SRRM2, we chose several mTORC1-associated genes (Supplementary Table S13, Supplementary Figure S7F) and compared the enzymatic activity of their encoded proteins after ArgRS or SRRM2 knock down. The enzyme Isocitrate dehydrogenase 1 (IDH1) responds to cell stress by regulating intracellular NADP/NADPH levels (Itsumi et al., 2015). Differential exon usage in *IDH1* correlated with an increase in IDH enzymatic activity in cell lysates upon SRRM2 knock down while ArgRS knock down led to decreased activity (Figure 7C). Differential exon usage was also found for genes encoding components of the ubiquitin/proteasome system (UPS), including the 19S proteasomal subunit de-ubiquitinating enzyme UCHL5 and the catalytically active proteasomal subunit PSMB5 (β5). These correlated with increased β5 activity upon SRRM2 knock down and decreased β5 activity upon ArgRS knock down (Figure 7D).

The UPS provides peptides that are presented on the cell surface by the major histocompatibility complex I (MHCI) and alternative splicing of UPS components, such as the β5 proteasomal subunit, could result in differences in peptide processing and consequentially peptides presented to immune cells by MHCI. Additionally, other splicing changes due to ArgRS and SRRM2 knock down that result in different protein isoforms could yield alternative peptides directly through differences in amino acid sequence and alternative protease cleavage sites. These MHCI-bound peptides communicate the status of the presenting cell to immune cells (Rock et al., 2016).

To investigate whether communication to immune cells through MHCI might be changed by ArgRS and SRRM2 knock down (Figure 7E), we compared their MHCI-bound peptidome using mass spectrometry. The detected MHCI-bound peptides matched previously described MHCI-bound peptides in length and a preference for the amino acids Tyr and Leu in position 2 (Bassani-Sternberg et al., 2015) (Supplementary Figure S7G). Overall, MHCI protein levels and the spectral counts for the measured peptides were comparable between all treatments (Supplementary Figure S7G, Figure 7F). However, upon sequencing of the peptides presented by MHCI by mass spectrometry, we found that ArgRS knock down substantially reduced the peptide diversity displayed on MHCI (20 vs 104). In contrast, SRRM2 knock down led to the presentation of a comparable number of different peptides by MHCI as the shSCR control (81 vs 104, Figure 7F, see also Supplementary Figure S7H). Nineteen of these peptides were unique for the SRRM2 knock down cell line and 41 were unique for shSCR control, suggesting that SRRM2 knock down led to the presentation of an altered set of MHCI-bound peptides (Figure 7F, Supplementary Figure S7H).

In order to predict how these MHCI peptide pools might influence communication with immune cells, we calculated MHCI immunogenicity scores (Figure 7F). A higher score predicts that these peptides would be more likely recognized by T cells and therefore more likely to elicit an immune response (Calis et al., 2013). Consistent with our hypothesis that ArgRS knock down mimics inflammation, MHCI peptides presented by ArgRS knock down cells had a higher median score than peptides that were presented only by SRRM2 knock down cells (Figure 7F). Collectively, these findings support a model in which alternative splicing events that are inversely regulated by ArgRS and SRRM2 could contribute to changes in cellular metabolism and communication as part of the cellular response to inflammation (Figure 7G).

## Discussion

Here we found that a portion of ArgRS localizes to the cell nucleus and that nuclear ArgRS levels fluctuate in an arginine-dependent manner (Figure 1). To characterize the function of ArgRS in the nucleus, we assessed the ArgRS interactome by mass spectrometry, identified SRRM2 as significantly and reproducibly enriched, and confirmed the ArgRS/SRRM2 interaction by reciprocal co-immunoprecipitation (Figure 2). Further, SRRM2 and ArgRS localized in proximity to each other in the nucleus (Figure 3). We showed that both arginine starvation and depletion of ArgRS increased SRRM2 mobility (Figure 4), suggesting that the interaction of nuclear ArgRS with SRRM2 might impede SRRM2 trafficking. Upon ArgRS knock down, depletion of nuclear ArgRS and altered SRRM2 mobility correlated with numerous changes in SRRM2-dependent alternative splicing, while small noncoding RNAs were largely unaffected by the ArgRS knock down (Figure 5, 6). We found further that SRRM2 and ArgRS knock downs modulated alternative splicing of a subset of genes in opposite directions, suggesting that ArgRS could affect SRRM2’s function (Figure 6). Two inversely regulated genes, *IDH1*, which encodes isocitrate dehydrogenase 1, and *PSMB5*, which encodes the β5 subunit of the 20S proteasome, showed that the inversely regulated alternative splicing changes had opposite effects on the biochemical activities of the encoded proteins. These changes in enzymatic activity impacted cellular metabolism and contributed to altered MHCI peptide presentation in a manner consistent with a cellular response to inflammation (Figure 7).

Arginine levels decrease during inflammation as pro- and anti-inflammatory enzymes compete for arginine as a substrate (Bronte and Zanovello, 2005). Eventually, the depletion of arginine leads to the resolution of inflammation due to a restraining effect on immune cells, especially T cells (Geiger et al., 2016; Murray, 2016). Therefore, a possible ArgRS/SRRM2 interaction and its consequences could have alternative outcomes at different stages of inflammation. Additionally, arginine is a key regulator of the mTOR pathway (Saxton et al., 2016; Wyant et al., 2017), and we found that expression of functionally diverse splice variants of mTORC1 pathway-associated genes (Figure 7B) - driven by the ArgRS and/or SRRM2 - had consequences for multiple functions of that pathway. Because manipulation of arginine triggers several cross-talking cellular signaling pathways (Bar-Peled and Sabatini, 2014; Dong et al., 2000), we focused on using ArgRS knock downs to specifically study ArgRS’ relationship with SRRM2. The observed downregulation of IDH1 activity and thereby NADPH production by ArgRS knock down for example would support the metabolic shutdown initiated by mTOR inhibition upon arginine depletion.

We showed previously that ArgRS’ presence in the nucleus is dependent on its presence in the MSC as nuclear ArgRS was strongly reduced upon exclusion of ArgRS from the MSC (Cui et al., 2021). Here, we found that ArgRS – once in the nucleus – interacted with SRRM2 independently of the MSC, as affinity enrichment of ArgRS, but not MetRS, another component of the MSC, retrieved SRRM2 (Figure 2B, lower right). These finding suggest that in addition to tRNA quality control (Lund and Dahlberg, 1998), the nuclear MSC might act as a reservoir of nuclear aaRSs with regulatory functions that are sequestered until a cue induces their release. This concept is similar to that suggested for the cytoplasmic MSC (Ray et al., 2007). We hypothesize that the nuclear MSC is a dynamic reservoir in steady exchange with its surroundings that enable the correct localization and the controlled release of their components. ArgRS, with changing arginine levels as a cue, thus connects two distinct protein organizational structures with central roles in transcription and translation. Due to their ability to sense and respond to changes in amino acids levels and interact with a variety of other proteins, aaRSs are uniquely poised to function in metabolite sensing and signal transduction (Netzer et al., 2009; Sajish and Schimmel, 2015; Son et al., 2019).

SRRM2 is a nuclear speckle protein and spliceosome component, and RNA transcription splicing have been reported to take place at or in close proximity to nuclear speckles (Belmont, 2021; Chen and Belmont, 2019). A recent model suggested that splice-site selection is regulated by partial immersion of unspliced pre-mRNA at the interface between nuclear speckles and nucleoplasm (Liao and Regev, 2021). The interaction between ArgRS and SRRM2 at the interface of SRRM2-rich areas could therefore impact alternative mRNA splicing directly by disrupting SRRM2’s function within the spliceosome and/or by regulating trafficking of SRRM2 to and from the nuclear speckle-nucleoplasm interface. Our interactome study would identify both direct and indirect interaction partners of ArgRS, as shown by the retrieval of MSC components that are not in direct contact with ArgRS. Therefore, we cannot exclude that additional biomolecules might be involved in mediating the physical interaction between ArgRS and SRRM2 or the effects observed upon ArgRS modulation. While we see accumulation of ArgRS/SRRM2 signal at the edge of SRRM2-rich regions, we cannot exclude that these proteins also interact outside of discrete nuclear structures, which would be more challenging to visualize by microscopy. The finding that the ArgRS/SRRM2 interaction could be retrieved by co-immunoprecipitation from cell lysates indicates that phase-separation is not a pre-requisite for binding.

SRRM2 is both essential for nuclear speckle formation (Ilik et al., 2020) and a component of active spliceosomes (Bertram et al., 2020). The N-terminal 1% of SRRM2 is immobilized sufficiently in the spliceosome to be resolved by cryo-EM while the remaining 99% (including the low complexity Ser/Arg-rich tail) are disordered (Bertram et al., 2020). SRRM2 would therefore be an ideal candidate to anchor and organize spliceosomes within or on the periphery of nuclear speckles. Due to its large size and many phosphorylation sites, SRRM2 provides ample interaction surface for other splicing regulatory proteins to gather in the proximity of the spliceosome by interacting with the exposed areas of SRRM2. ArgRS could impede SRRM2’s function as a splicing factor either by preventing interaction with other splicesomal proteins - including its integration into the spliceosome - or by hindering SRRM2 mobility. More generally, our results suggest that SRRM2 function could be regulated by its trafficking in the nucleus and raise the possibility that other regulatory proteins could function by modulating this trafficking comparable to ArgRS.

A previous study in a murine liver cell line found that depletion of SRRM2 to lower levels than in our study resulted in upregulation of transcription and splicing of immune response genes (Hu et al., 2019). Here, we found that modulation of SRRM2 levels impacted cellular metabolism and MCHI peptide processing (Figure 7). MHCI presentation allows cells to communicate with the immune system (Rock et al., 2016). In pathologies, such as viral infection and cancer, an altered MHCI peptidome could induce different immune responses. We found that ArgRS knock down, which mimics arginine starvation by leading to decreased concentrations of ArgRS in the nucleus, led to a decrease in peptide diversity without reducing the total amount of detectable MHCI-presented peptides (Figure 7F).

These peptides were also predicted to be more immunogenic (Figure 7F) and might therefore be preferentially recognized for example by cytotoxic T cells during cancer and viral infections, in which inflammation is induced. Under inflammatory conditions, the reduction of peptide diversity and increase in immunogenicity presented by MHCI upon ArgRS knock down could enable more efficient identification by the immune system. On the other hand, SRRM2 knock down led to the presentation of a different set of MHCI peptides as compared to control (Figure 7F), suggesting that sequestration of SRRM2 by ArgRS could potentially increase the diversity of MHCI peptide presentation. Those MHCI peptides exclusively found on SRRM2 knock down cells were predicted to be less immunogenic (Figure 7F). Recently, splicing inhibitors have been used to modulate the MHCI peptidome to increase sensitivity to immune therapy in cancer (Lu et al., 2021) and our findings suggest that a similar mechanism is used to modulate and diversify MHCI peptide variety is employed during inflammation. The interplay found here between arginine, ArgRS, and SRRM2 would enable cells to not only detect ongoing extracellular inflammation by picking up metabolic cues but also to respond adequately by adapting their proteome and tuning their communication with immune cells.

## Supporting information

Resource Table

## Acknowledgements

We thank the Center for Metabolomics, the Microscopy core, the High Performance Computing team, and Flow Cytometry Core, all at Scripps Research, for training and instrument maintenance, and the staff at the Genomic Sequencing and Analysis Facility at the University of Texas at Austin for poly(A) library preparation and sequencing. Access to a plate reader and flow cytometry software was provided by Prof. James Paulson and access to a ChemiDoc Imager by Prof. Jeffrey Kelly, both at Scripps Research. We thank Aditi Dutta for preliminary analysis of ArgRS- and MetRS-dependent signaling pathways. We also thank Dr. Ulrich Braunschweig, Dr. Rasim Barutcu, and Mingkun Wu (all University of Toronto) for feedback on the experimental design (UB) and the manuscript (RB, MW). We thank Professors David Sabatini, Phillip A. Sharp, and Landon J. Edgar for reading the manuscript and their insightful comments.

## Author contributions

HC planned and conducted experiments and analyzed data. JKD, JJM, and JRY performed, analyzed, or supervised mass spectrometry experiments. DCW, RMN, and AML performed, analyzed, or supervised TGIRT-seq and RNA-seq library preparation and sequencing. JJL and BB analyzed RNA-seq results. PS, AML, and HC designed research and wrote the manuscript with input and contributions from all authors.

## Data and materials availability

TGIRT-seq data are deposited in the NCBI sequence read archive and accessible through the BioProject number PRJNA561913. Poly(A) RNA-seq data are deposited GEO GSE165513. Mass spectrometry data is deposited on ProteomeXchange Consortium via the PRIDE partner repository with the dataset identifiers PXD015692 (interactomes), PXD024091 (N-terminomics), and PXD027531 (MHCI peptidome). All unique reagents generated in this study are available from Prof. Paul Schimmel, with a completed Materials Transfer Agreement.

## List of Supplementary materials

Material and Methods

Supplementary Figures S1-S7

Supplementary Tables S1-S15

Supplementary Videos S1-S7

## References

Aass, K.R., Kastnes, M.H., and Standal, T. (2021). Molecular interactions and functions of IL-32. J. Leukoc. Biol. 109, 143–159.

Anders, S., Reyes, A., and Huber, W. (2012). Detecting differential usage of exons from RNA-seq data. Genome Res. 22, 2008–2017.

Bar-Peled, L., and Sabatini, D.M. (2014). Regulation of mTORC1 by amino acids. Trends Cell Biol. 24, 400–406.

Bassani-Sternberg, M., Pletscher-Frankild, S., Jensen, L.J., and Mann, M. (2015). Mass spectrometry of human leukocyte antigen class I peptidomes reveals strong effects of protein abundance and turnover on antigen presentation. Mol. Cell. Proteomics MCP 14, 658–673.

Belmont, A.S. (2021). Nuclear Compartments: An Incomplete Primer to Nuclear Compartments, Bodies, and Genome Organization Relative to Nuclear Architecture. Cold Spring Harb. Perspect. Biol. a041268.

Bertram, K., El Ayoubi, L., Dybkov, O., Agafonov, D.E., Will, C.L., Hartmuth, K., Urlaub, H., Kastner, B., Stark, H., and Lührmann, R. (2020). Structural Insights into the Roles of Metazoan-Specific Splicing Factors in the Human Step 1 Spliceosome. Mol. Cell 80, 127–139.e6.

Blencowe, B.J. (2017). The Relationship between Alternative Splicing and Proteomic Complexity. Trends Biochem. Sci. 42, 407–408.

Blencowe, B.J., Issner, R., Nickerson, J.A., and Sharp, P.A. (1998). A coactivator of pre-mRNA splicing. Genes Dev. 12, 996–1009.

Blencowe, B.J., Baurén, G., Eldridge, A.G., Issner, R., Nickerson, J.A., Rosonina, E., and Sharp, P.A. (2000). The SRm160/300 splicing coactivator subunits. RNA N. Y. N 6, 111–120.

Boivin, V., Deschamps-Francoeur, G., Couture, S., Nottingham, R.M., Bouchard-Bourelle, P., Lambowitz, A.M., Scott, M.S., and Abou-Elela, S. (2018). Simultaneous sequencing of coding and noncoding RNA reveals a human transcriptome dominated by a small number of highly expressed noncoding genes. RNA N. Y. N 24, 950–965.

Brangwynne, C.P., Eckmann, C.R., Courson, D.S., Rybarska, A., Hoege, C., Gharakhani, J., Jülicher, F., and Hyman, A.A. (2009). Germline P granules are liquid droplets that localize by controlled dissolution/condensation. Science 324, 1729–1732.

Braunschweig, U., Barbosa-Morais, N.L., Pan, Q., Nachman, E.N., Alipanahi, B., Gonatopoulos-Pournatzis, T., Frey, B., Irimia, M., and Blencowe, B.J. (2014). Widespread intron retention in mammals functionally tunes transcriptomes. Genome Res. 24, 1774–1786.

Bronte, V., and Zanovello, P. (2005). Regulation of immune responses by L-arginine metabolism. Nat. Rev. Immunol. 5, 641–654.

Calis, J.J.A., Maybeno, M., Greenbaum, J.A., Weiskopf, D., De Silva, A.D., Sette, A., Keşmir, C., and Peters, B. (2013). Properties of MHC class I presented peptides that enhance immunogenicity. PLoS Comput. Biol. 9, e1003266.

Chen, Y., and Belmont, A.S. (2019). Genome organization around nuclear speckles. Curr. Opin. Genet. Dev. 55, 91–99.

Costes, S.V., Daelemans, D., Cho, E.H., Dobbin, Z., Pavlakis, G., and Lockett, S. (2004). Automatic and quantitative measurement of protein-protein colocalization in live cells. Biophys. J. 86, 3993–4003.

Cui, H., Kapur, M., Diedrich, J.K., Yates, J.R., Ackerman, S.L., and Schimmel, P. (2021). Regulation of ex-translational activities is the primary function of the multi-tRNA synthetase complex. Nucleic Acids Res. 49, 3603–3616.

Dong, J., Qiu, H., Garcia-Barrio, M., Anderson, J., and Hinnebusch, A.G. (2000). Uncharged tRNA activates GCN2 by displacing the protein kinase moiety from a bipartite tRNA-binding domain. Mol. Cell 6, 269–279.

Duchon, A.A., St Gelais, C., Titkemeier, N., Hatterschide, J., Wu, L., and Musier-Forsyth, K. (2017). HIV-1 Exploits a Dynamic Multi-aminoacyl-tRNA Synthetase Complex To Enhance Viral Replication. J. Virol. 91.

Galarza-Muñoz, G., Briggs, F.B.S., Evsyukova, I., Schott-Lerner, G., Kennedy, E.M., Nyanhete, T., Wang, L., Bergamaschi, L., Widen, S.G., Tomaras, G.D., et al. (2017). Human Epistatic Interaction Controls IL7R Splicing and Increases Multiple Sclerosis Risk. Cell 169, 72–84.e13.

Galganski, L., Urbanek, M.O., and Krzyzosiak, W.J. (2017). Nuclear speckles: molecular organization, biological function and role in disease. Nucleic Acids Res. 45, 10350–10368.

Gautam, A., Grainger, R.J., Vilardell, J., Barrass, J.D., and Beggs, J.D. (2015). Cwc21p promotes the second step conformation of the spliceosome and modulates 3’ splice site selection. Nucleic Acids Res. 43, 3309–3317.

Geiger, R., Rieckmann, J.C., Wolf, T., Basso, C., Feng, Y., Fuhrer, T., Kogadeeva, M., Picotti, P., Meissner, F., Mann, M., et al. (2016). L-Arginine Modulates T Cell Metabolism and Enhances Survival and Anti-tumor Activity. Cell 167, 829–842.e13.

Guo, M., and Schimmel, P. (2013). Essential nontranslational functions of tRNA synthetases. Nat. Chem. Biol. 9, 145–153.

Guo, Y.E., Manteiga, J.C., Henninger, J.E., Sabari, B.R., Dall’Agnese, A., Hannett, N.M., Spille, J.-H., Afeyan, L.K., Zamudio, A.V., Shrinivas, K., et al. (2019). Pol II phosphorylation regulates a switch between transcriptional and splicing condensates. Nature 572, 543–548.

Han, H., Braunschweig, U., Gonatopoulos-Pournatzis, T., Weatheritt, R.J., Hirsch, C.L., Ha, K.C.H., Radovani, E., Nabeel-Shah, S., Sterne-Weiler, T., Wang, J., et al. (2017). Multilayered Control of Alternative Splicing Regulatory Networks by Transcription Factors. Mol. Cell 65, 539–553.e7.

Hayano, M., Yang, W.S., Corn, C.K., Pagano, N.C., and Stockwell, B.R. (2016). Loss of cysteinyl-tRNA synthetase (CARS) induces the transsulfuration pathway and inhibits ferroptosis induced by cystine deprivation. Cell Death Differ. 23, 270–278.

Hnisz, D., Shrinivas, K., Young, R.A., Chakraborty, A.K., and Sharp, P.A. (2017). A Phase Separation Model for Transcriptional Control. Cell 169, 13–23.

Horiguchi, N., Lafdil, F., Miller, A.M., Park, O., Wang, H., Rajesh, M., Mukhopadhyay, P., Fu, X.Y., Pacher, P., and Gao, B. (2010). Dissociation between liver inflammation and hepatocellular damage induced by carbon tetrachloride in myeloid cell-specific signal transducer and activator of transcription 3 gene knockout mice. Hepatol. Baltim. Md 51, 1724–1734.

Hu, S., Lv, P., Yan, Z., and Wen, B. (2019). Disruption of nuclear speckles reduces chromatin interactions in active compartments. Epigenetics Chromatin 12, 43.

Ilik, İ.A., Malszycki, M., Lübke, A.K., Schade, C., Meierhofer, D., and Aktaş, T. (2020). SON and SRRM2 are essential for nuclear speckle formation. ELife 9, e60579.

Irimia, M., Weatheritt, R.J., Ellis, J.D., Parikshak, N.N., Gonatopoulos-Pournatzis, T., Babor, M., Quesnel-Vallières, M., Tapial, J., Raj, B., O’Hanlon, D., et al. (2014). A highly conserved program of neuronal microexons is misregulated in autistic brains. Cell 159, 1511–1523.

Itsumi, M., Inoue, S., Elia, A.J., Murakami, K., Sasaki, M., Lind, E.F., Brenner, D., Harris, I.S., Chio, I.I.C., Afzal, S., et al. (2015). Idh1 protects murine hepatocytes from endotoxin-induced oxidative stress by regulating the intracellular NADP(+)/NADPH ratio. Cell Death Differ. 22, 1837–1845.

Jones, R.G., and Pearce, E.J. (2017). MenTORing Immunity: mTOR Signaling in the Development and Function of Tissue-Resident Immune Cells. Immunity 46, 730–742.

Kaminska, M., Havrylenko, S., Decottignies, P., Gillet, S., Le Maréchal, P., Negrutskii, B., and Mirande, M. (2009). Dissection of the structural organization of the aminoacyl-tRNA synthetase complex. J. Biol. Chem. 284, 6053–6060.

Kaplanis, J., Samocha, K.E., Wiel, L., Zhang, Z., Arvai, K.J., Eberhardt, R.Y., Gallone, G., Lelieveld, S.H., Martin, H.C., McRae, J.F., et al. (2020). Evidence for 28 genetic disorders discovered by combining healthcare and research data. Nature 1–7.

Keilhauer, E.C., Hein, M.Y., and Mann, M. (2015). Accurate protein complex retrieval by affinity enrichment mass spectrometry (AE-MS) rather than affinity purification mass spectrometry (AP-MS). Mol. Cell. Proteomics MCP 14, 120–135.

Kim, M.H., and Kim, S. (2020). Chapter Six - Structures and functions of multi-tRNA synthetase complexes. In The Enzymes, L. Ribas de Pouplana, and L.S. Kaguni, eds. (Academic Press), pp. 149–173.

Kleifeld, O., Doucet, A., Prudova, A., auf dem Keller, U., Gioia, M., Kizhakkedathu, J.N., and Overall, C.M. (2011). Identifying and quantifying proteolytic events and the natural N terminome by terminal amine isotopic labeling of substrates. Nat. Protoc. 6, 1578–1611.

Kwon, N.H., Fox, P.L., and Kim, S. (2019). Aminoacyl-tRNA synthetases as therapeutic targets. Nat. Rev. Drug Discov. 18, 629–650.

Lee, S.W., Cho, B.H., Park, S.G., and Kim, S. (2004). Aminoacyl-tRNA synthetase complexes: beyond translation. J. Cell Sci. 117, 3725–3734.

Liao, S.E., and Regev, O. (2021). Splicing at the phase-separated nuclear speckle interface: a model. Nucleic Acids Res. 49, 636–645.

Lu, S.X., De Neef, E., Thomas, J.D., Sabio, E., Rousseau, B., Gigoux, M., Knorr, D.A., Greenbaum, B., Elhanati, Y., Hogg, S.J., et al. (2021). Pharmacologic modulation of RNA splicing enhances anti-tumor immunity. Cell 184, 4032–4047.e31.

Lund, E., and Dahlberg, J.E. (1998). Proofreading and aminoacylation of tRNAs before export from the nucleus. Science 282, 2082–2085.

Miyagawa, R., Tano, K., Mizuno, R., Nakamura, Y., Ijiri, K., Rakwal, R., Shibato, J., Masuo, Y., Mayeda, A., Hirose, T., et al. (2012). Identification of cis- and trans-acting factors involved in the localization of MALAT-1 noncoding RNA to nuclear speckles. RNA N. Y. N 18, 738–751.

Murray, P.J. (2016). Amino acid auxotrophy as a system of immunological control nodes. Nat. Immunol. 17, 132–139.

Nathanson, L., and Deutscher, M.P. (2000). Active aminoacyl-tRNA synthetases are present in nuclei as a high molecular weight multienzyme complex. J. Biol. Chem. 275, 31559–31562.

Netea, M.G., Balkwill, F., Chonchol, M., Cominelli, F., Donath, M.Y., Giamarellos-Bourboulis, E.J., Golenbock, D., Gresnigt, M.S., Heneka, M.T., Hoffman, H.M., et al. (2017). A guiding map for inflammation. Nat. Immunol. 18, 826–831.

Netzer, N., Goodenbour, J.M., David, A., Dittmar, K.A., Jones, R.B., Schneider, J.R., Boone, D., Eves, E.M., Rosner, M.R., Gibbs, J.S., et al. (2009). Innate immune and chemically triggered oxidative stress modifies translational fidelity. Nature 462, 522–526.

Nottingham, R.M., Wu, D.C., Qin, Y., Yao, J., Hunicke-Smith, S., and Lambowitz, A.M. (2016). RNA-seq of human reference RNA samples using a thermostable group II intron reverse transcriptase. RNA N. Y. N 22, 597–613.

Ofir-Birin, Y., Fang, P., Bennett, S.P., Zhang, H.-M., Wang, J., Rachmin, I., Shapiro, R., Song, J., Dagan, A., Pozo, J., et al. (2013). Structural switch of lysyl-tRNA synthetase between translation and transcription. Mol. Cell 49, 30–42.

Pan, Q., Shai, O., Misquitta, C., Zhang, W., Saltzman, A.L., Mohammad, N., Babak, T., Siu, H., Hughes, T.R., Morris, Q.D., et al. (2004). Revealing global regulatory features of mammalian alternative splicing using a quantitative microarray platform. Mol. Cell 16, 929–941.

Patil, M.D., Bhaumik, J., Babykutty, S., Banerjee, U.C., and Fukumura, D. (2016). Arginine dependence of tumor cells: targeting a chink in cancer’s armor. Oncogene 35, 4957–4972.

Phair, R.D., and Misteli, T. (2000). High mobility of proteins in the mammalian cell nucleus. Nature 404, 604–609.

Qin, Y., Yao, J., Wu, D.C., Nottingham, R.M., Mohr, S., Hunicke-Smith, S., and Lambowitz, A.M. (2016). High-throughput sequencing of human plasma RNA by using thermostable group II intron reverse transcriptases. RNA N. Y. N 22, 111–128.

Ray, P.S., Arif, A., and Fox, P.L. (2007). Macromolecular complexes as depots for releasable regulatory proteins. Trends Biochem. Sci. 32, 158–164.

Rock, K.L., Reits, E., and Neefjes, J. (2016). Present Yourself! By MHC Class I and MHC Class II Molecules. Trends Immunol. 37, 724–737.

Saitoh, N., Spahr, C.S., Patterson, S.D., Bubulya, P., Neuwald, A.F., and Spector, D.L. (2004). Proteomic analysis of interchromatin granule clusters. Mol. Biol. Cell 15, 3876–3890.

Sajish, M., and Schimmel, P. (2015). A human tRNA synthetase is a potent PARP1-activating effector target for resveratrol. Nature 519, 370–373.

Sarkar, S., Azad, A.K., and Hopper, A.K. (1999). Nuclear tRNA aminoacylation and its role in nuclear export of endogenous tRNAs in Saccharomyces cerevisiae. Proc. Natl. Acad. Sci. U. S. A. 96, 14366–14371.

Saxton, R.A., Chantranupong, L., Knockenhauer, K.E., Schwartz, T.U., and Sabatini, D.M. (2016). Mechanism of arginine sensing by CASTOR1 upstream of mTORC1. Nature 536, 229–233.

Schimmel, P.R., and Söll, D. (1979). Aminoacyl-tRNA synthetases: general features and recognition of transfer RNAs. Annu. Rev. Biochem. 48, 601–648.

Seburn, K.L., Nangle, L.A., Cox, G.A., Schimmel, P., and Burgess, R.W. (2006). An active dominant mutation of glycyl-tRNA synthetase causes neuropathy in a Charcot-Marie-Tooth 2D mouse model. Neuron 51, 715–726.

Shehadeh, L.A., Yu, K., Wang, L., Guevara, A., Singer, C., Vance, J., and Papapetropoulos, S. (2010). SRRM2, a potential blood biomarker revealing high alternative splicing in Parkinson’s disease. PloS One 5, e9104.

Shi, Y., Xu, X., Zhang, Q., Fu, G., Mo, Z., Wang, G.S., Kishi, S., and Yang, X.-L. (2014). tRNA synthetase counteracts c-Myc to develop functional vasculature. ELife 3, e02349.

Shi, Y., Wei, N., and Yang, X.-L. (2017). Studying nuclear functions of aminoacyl tRNA synthetases. Methods San Diego Calif 113, 105–110.

Shin, Y., and Brangwynne, C.P. (2017). Liquid phase condensation in cell physiology and disease. Science 357.

Smith, K.P., Hall, L.L., and Lawrence, J.B. (2020). Nuclear hubs built on RNAs and clustered organization of the genome. Curr. Opin. Cell Biol. 64, 67–76.

Son, K., You, J.-S., Yoon, M.-S., Dai, C., Kim, J.H., Khanna, N., Banerjee, A., Martinis, S.A., Han, G., Han, J.M., et al. (2019). Nontranslational function of leucyl-tRNA synthetase regulates myogenic differentiation and skeletal muscle regeneration. J. Clin. Invest. 129, 2088–2093.

Spector, D.L., and Lamond, A.I. (2011). Nuclear speckles. Cold Spring Harb. Perspect. Biol. 3.

Tapial, J., Ha, K.C.H., Sterne-Weiler, T., Gohr, A., Braunschweig, U., Hermoso-Pulido, A., Quesnel-Vallières, M., Permanyer, J., Sodaei, R., Marquez, Y., et al. (2017). An atlas of alternative splicing profiles and functional associations reveals new regulatory programs and genes that simultaneously express multiple major isoforms. Genome Res. 27, 1759–1768.

Tomsic, J., He, H., Akagi, K., Liyanarachchi, S., Pan, Q., Bertani, B., Nagy, R., Symer, D.E., Blencowe, B.J., and de la Chapelle, A. (2015). A germline mutation in SRRM2, a splicing factor gene, is implicated in papillary thyroid carcinoma predisposition. Sci. Rep. 5, 10566.

Weichhart, T., Hengstschläger, M., and Linke, M. (2015). Regulation of innate immune cell function by mTOR. Nat. Rev. Immunol. 15, 599–614.

Wyant, G.A., Abu-Remaileh, M., Wolfson, R.L., Chen, W.W., Freinkman, E., Danai, L.V., Vander Heiden, M.G., and Sabatini, D.M. (2017). mTORC1 Activator SLC38A9 Is Required to Efflux Essential Amino Acids from Lysosomes and Use Protein as a Nutrient. Cell 171, 642–654.e12.

Yannay-Cohen, N., Carmi-Levy, I., Kay, G., Yang, C.M., Han, J.M., Kemeny, D.M., Kim, S., Nechushtan, H., and Razin, E. (2009). LysRS serves as a key signaling molecule in the immune response by regulating gene expression. Mol. Cell 34, 603–611.

Zanini, I.M.Y., Soneson, C., Lorenzi, L.E., and Azzalin, C.M. (2017). Human cactin interacts with DHX8 and SRRM2 to assure efficient pre-mRNA splicing and sister chromatid cohesion. J. Cell Sci. 130, 767–778.

Zhang, X., Yan, C., Hang, J., Finci, L.I., Lei, J., and Shi, Y. (2017). An Atomic Structure of the Human Spliceosome. Cell 169, 918–929.e14.

Zhu, L., and Brangwynne, C.P. (2015). Nuclear bodies: the emerging biophysics of nucleoplasmic phases. Curr. Opin. Cell Biol. 34, 23–30.

