## Supplementary Figures for "Arg-tRNA synthetase links inflammatory metabolism to RNA splicing and nuclear trafficking via SRRM2"

### Supplementary Figure S1

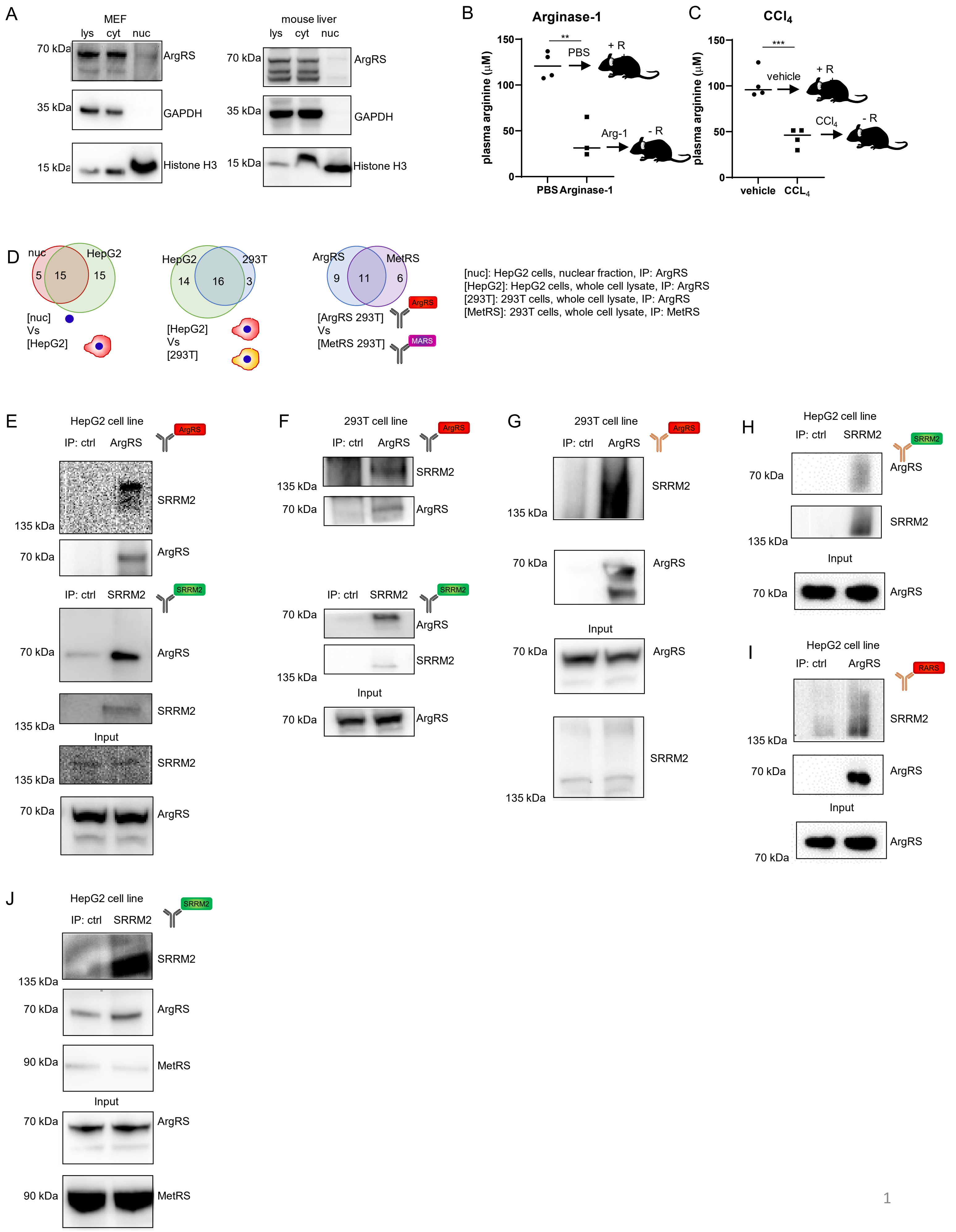

**Supplementary Figure 1: ArgRS localizes to the nucleus in murine cells and tissues, plasma arginine levels, and characterization of the ArgRS interactome.** (A) ArgRS in different cellular compartments of murine embryonic fibroblasts (MEF, left) and mouse liver (right) probed by western blot following cell fractionation. GAPDH: cytoplasmic marker, Histone H3: nuclear marker. lys: lysate, cyt: cytoplasm, nuc: nucleus. (B, C) LC/MS measurement of plasma arginine after (B) treatment with recombinant Arginase-1, an arginine-degrading enzyme, or (C) by induction of systemic inflammation with the liver toxin $CCl_4$. n=3 or 4 mice in each group. p= **0.002, p= *** p= 0.0009. (D) Venn diagrams comparing 4 different ArgRS interactomes. Statistical significance of overrepresentation by hypergeometric test: [nuc] vs [HepG2]: p=7.6e-10, [HepG2] vs [293T]: p=1.9e-10, [ArgRS 293T] vs [MetRS 293T]: p=2.0e-7. (E, F) SRRM2 interaction with ArgRS was verified by immunoprecipitation (IP) of ArgRS and vice versa. Detection of ArgRS and SRRM2 by western blot. Ctrl: Non-targeting antibody of the same isotype. (E) Input: HepG2 cell lysate. (F) Input: HEK 293T cell lysate. (G, H, I) Verification of ArgRS and SRRM2 interaction by immunoprecipitation (IP) with an alternative set of antibodies. (G) Input: 293T cell lysate. IP with ArgRS antibody. (H) Input: HepG2 cell lysate. IP with SRRM2 antibody. (I) HepG2 cell lysate. IP with ArgRS antibody. (J) SRRM2 immunoprecipitation did not retrieve MetRS but ArgRS. Detection of ArgRS, MetRS, and SRRM2 by western blot.

### Supplementary Figure S2

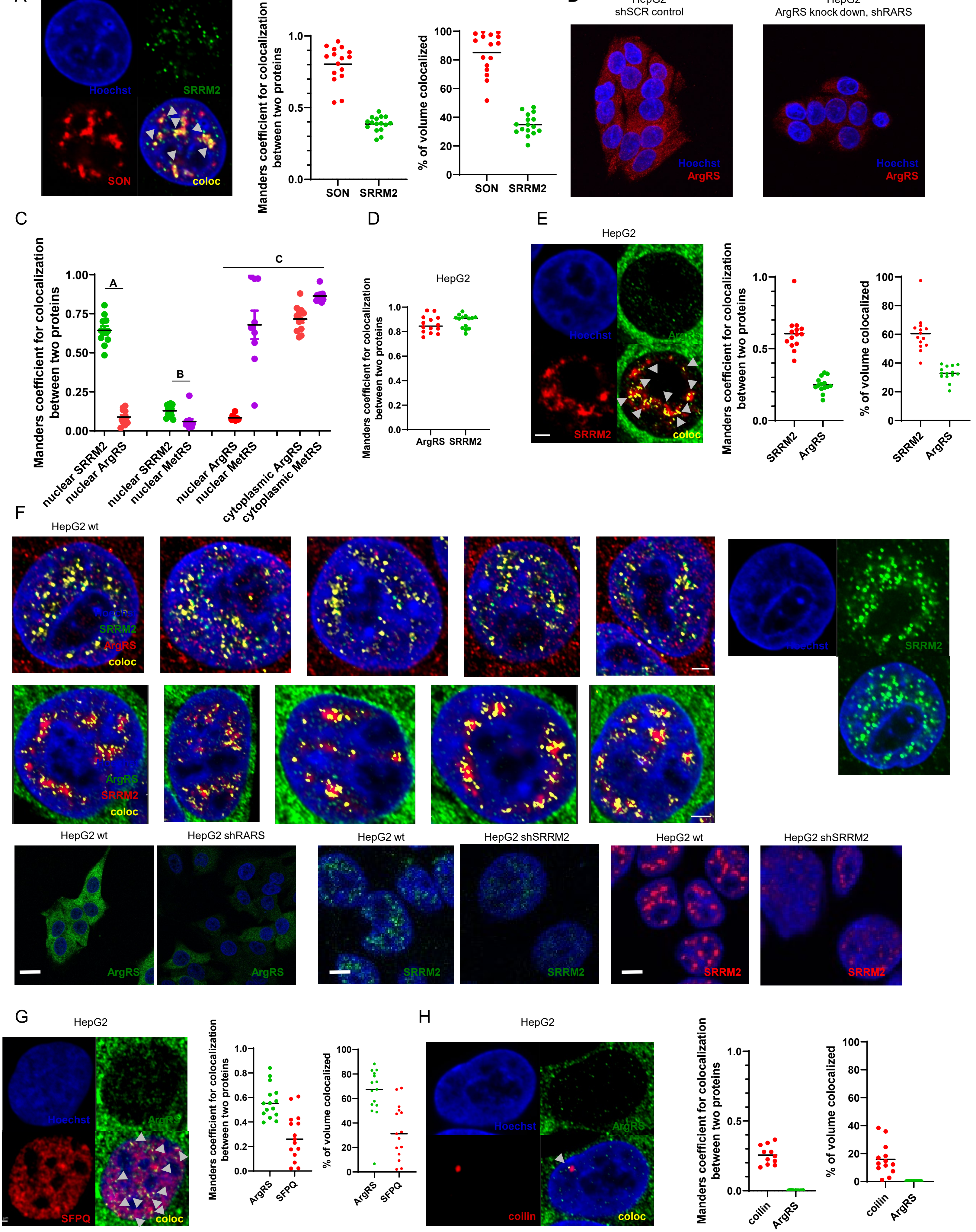

#### Supplementary Figure S2

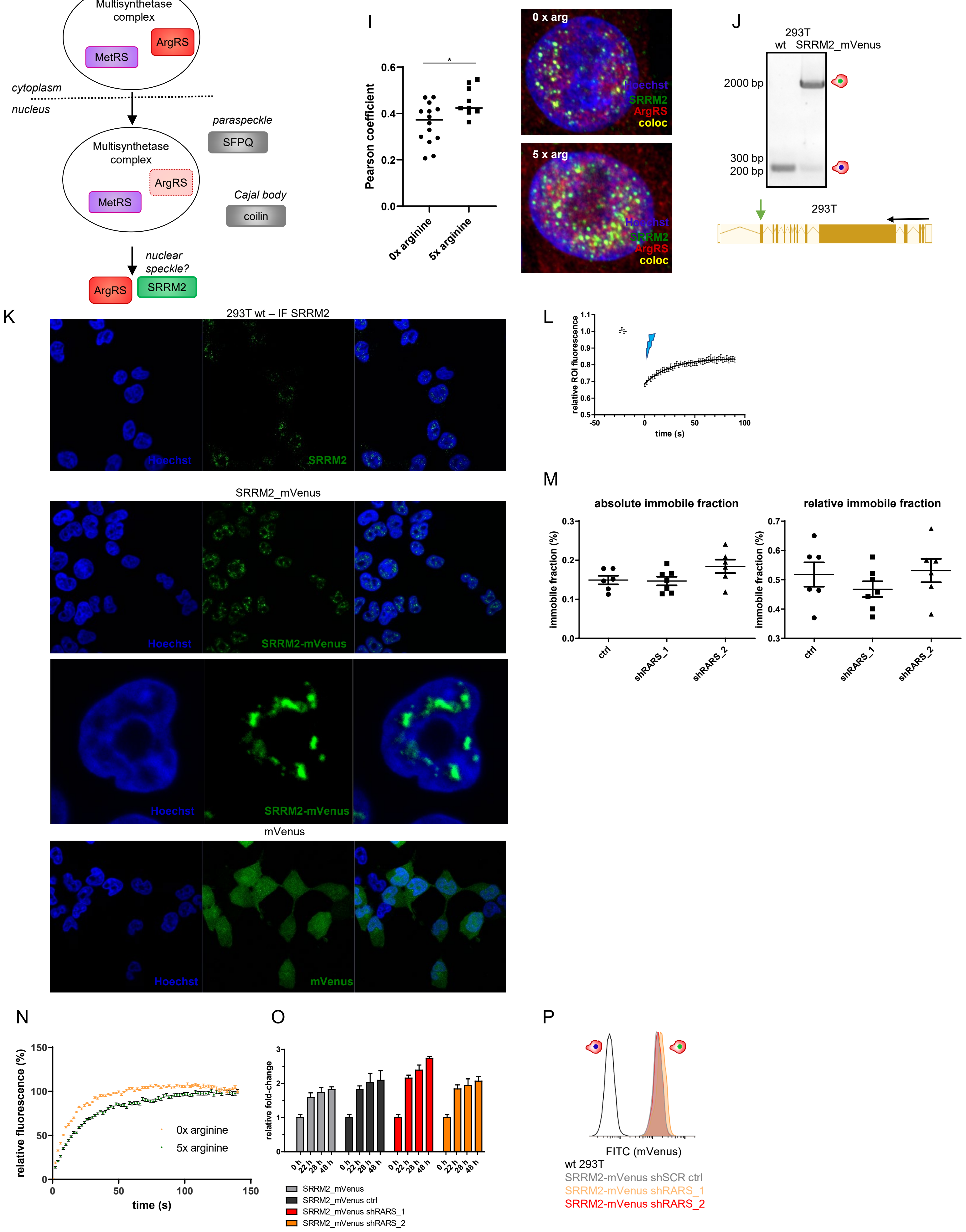

**Supplementary Figure 2: Fluorescent microscopy to assess SRRM2 colocalization and dynamics.** (A) Colocalization (yellow) of SRRM2 (green) and speckle protein SON (red) by immunofluorescent staining and imaging by confocal microscopy. 84% of SON colocalizes with SRRM2 and 34% of SRRM2 colocalizes with SON. (B) Maximum intensity projections of ArgRS immunofluorescent staining in HepG2 shSCR control and shRARS_1 cells to show antibody specificity. (C) Manders coefficients corresponding to panels A-C. Hoechst staining was used to identify the cell nucleus. Manders coefficient was calculated for individual cell nuclei (A, B, n = 11, 12) or for an average for 2-3 cells (C, n = 8). (D) Manders coefficient of ArgRS and SRRM2 colocalization when Costes thresholding was used. (E) Colocalization (yellow) of ArgRS (green) and SRRM2 (red) with different antibodies than used in Figure 2A. Manders coefficients of 14 cells. (F) Immunofluorescent staining of ArgRS and SRRM2 in representative nuclei and verification of antibodies by ArgRS and SRRM2 knock down. Bar: 2 µm. First row, right panel: SRRM2 staining without signal intensity threshold shows diffuse areas with weaker staining in the nucleus. Lower panel: verification of antibodies by ArgRS (shRARS) and SRRM2 (shSRRM2) knock down. Bar: wt and shRARS, 20 µm. wt and shSRRM2: 5 µm. (G) Colocalization of ArgRS (green) and paraspeckle protein SFPQ (red). Manders coefficient of colocalization after Costes thresholding. (H) Colocalization of ArgRS (green) and Cajal body protein coilin (red). Manders coefficient of colocalization. Scheme: Localization of proteins within different nuclear compartments. (I) Pearson correlation coefficient of SRRM2 and ArgRS colocalization by immunofluorescence staining upon arginine starvation (0x arginine) and high arginine (5x arginine). Representative immunofluorescent staining of ArgRS and SRRM2. p=*0.01, n=14, 9. Arginine concentration relative to DMEM. (J) Labeling of endogenous SRRM2 with the fluorescent protein mVenus. Verification of genomic mVenus insertion by PCR. Predicted amplification length of wildtype amplicon: 238 bp; predicted length of SRRM2-mVenus amplicon: 2071 bp. (K) Labeling with mVenus did not alter SRRM2 localization in 293T cells. Endogenous, unlabeled SRRM2 was visualized by immunofluorescence (upper panel). SRRM2-mVenus fluorescence displayed a similar pattern with fluorescence restricted to the cell nucleus and defined foci (middle panel). In comparison, stable expression of mVenus alone did not display a distinct localization (lower panel). (L) Fluorescence Recovery After Photobleaching (FRAP) measured over 90 s. Recovery reached its plateau after 60 s. (M) Absolute immobile SRRM2-mVenus fraction calculated from one-exponential curve fits of FRAP measurements and relative immobile fraction calculated by subtraction of residual fluorescence directly after bleaching. (N) Recovery of fluorescent signal is faster in the absence of arginine. Fluorescent signal was corrected for background and normalized. Each measurement included 14 regions of interest and the geometric mean of 8 measurements is shown. Errors bars depict standard error of the mean. (O) Cell viability upon ArgRS knock down in SRRM2-mVenus cells, measured with Alamar Blue. Technical quadruplets are shown. (P) Knock down of ArgRS did not affect SRRM2-mVenus protein levels measured by flow cytometry.

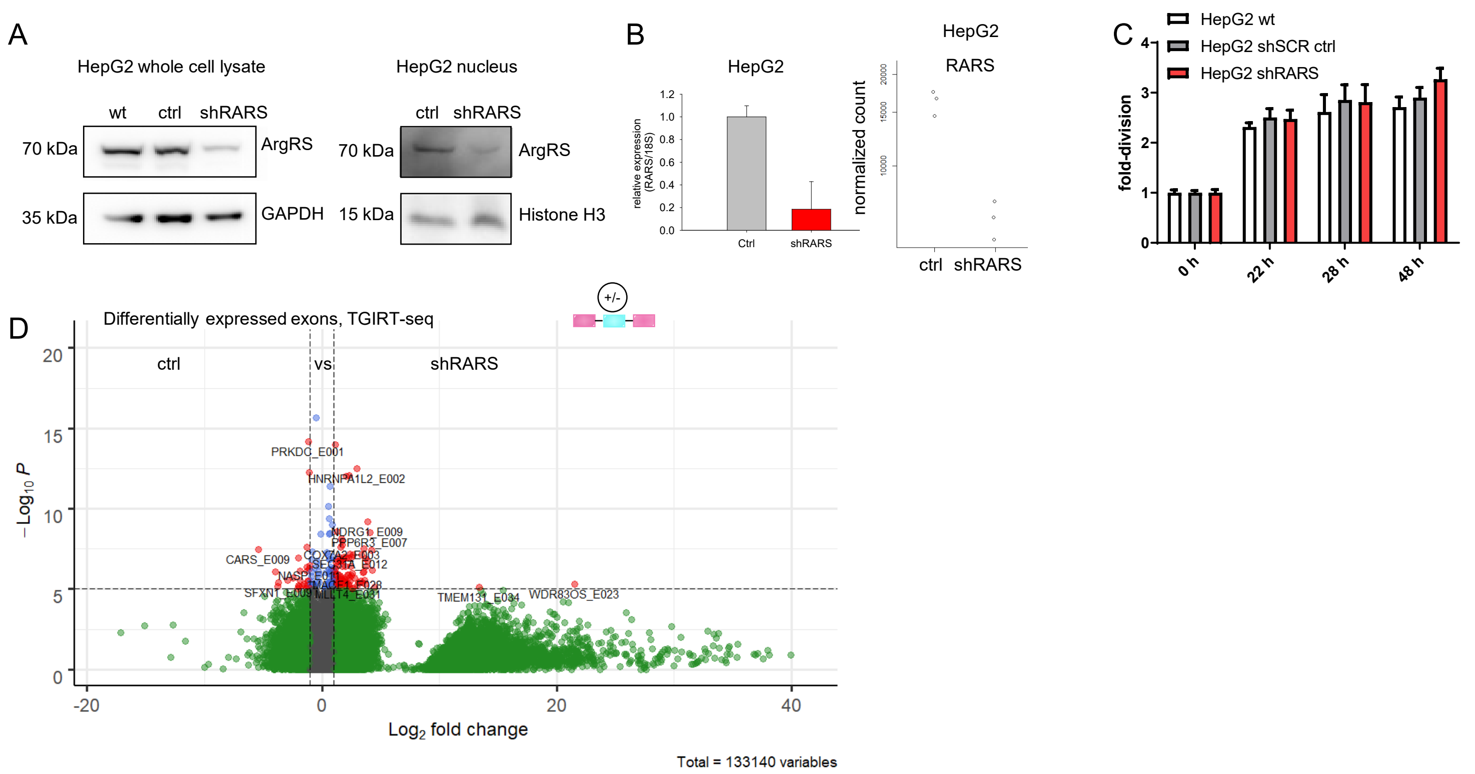


**Supplementary Figure 3: Characterization of ArgRS knock down.** (A) ArgRS knock down was confirmed on protein level by western blot and (B) on mRNA level by RT-qPCR and RNA-seq. A representative out of 3 repeats is shown. (C) Cell viability and proliferation of HepG2 cells was not affected by ArgRS knock down. Alamar blue based cell viability assay, quadruplets of technical replicates are shown. (D) Volcano plot of differentially expressed exons upon ArgRS knock down as found by TGIRT-seq. Dashed vertical lines: 2-fold change, dashed horizontal line: p-value 1e-6. (A-D) ctrl: shSCR control, shRARS: ArgRS knock down.

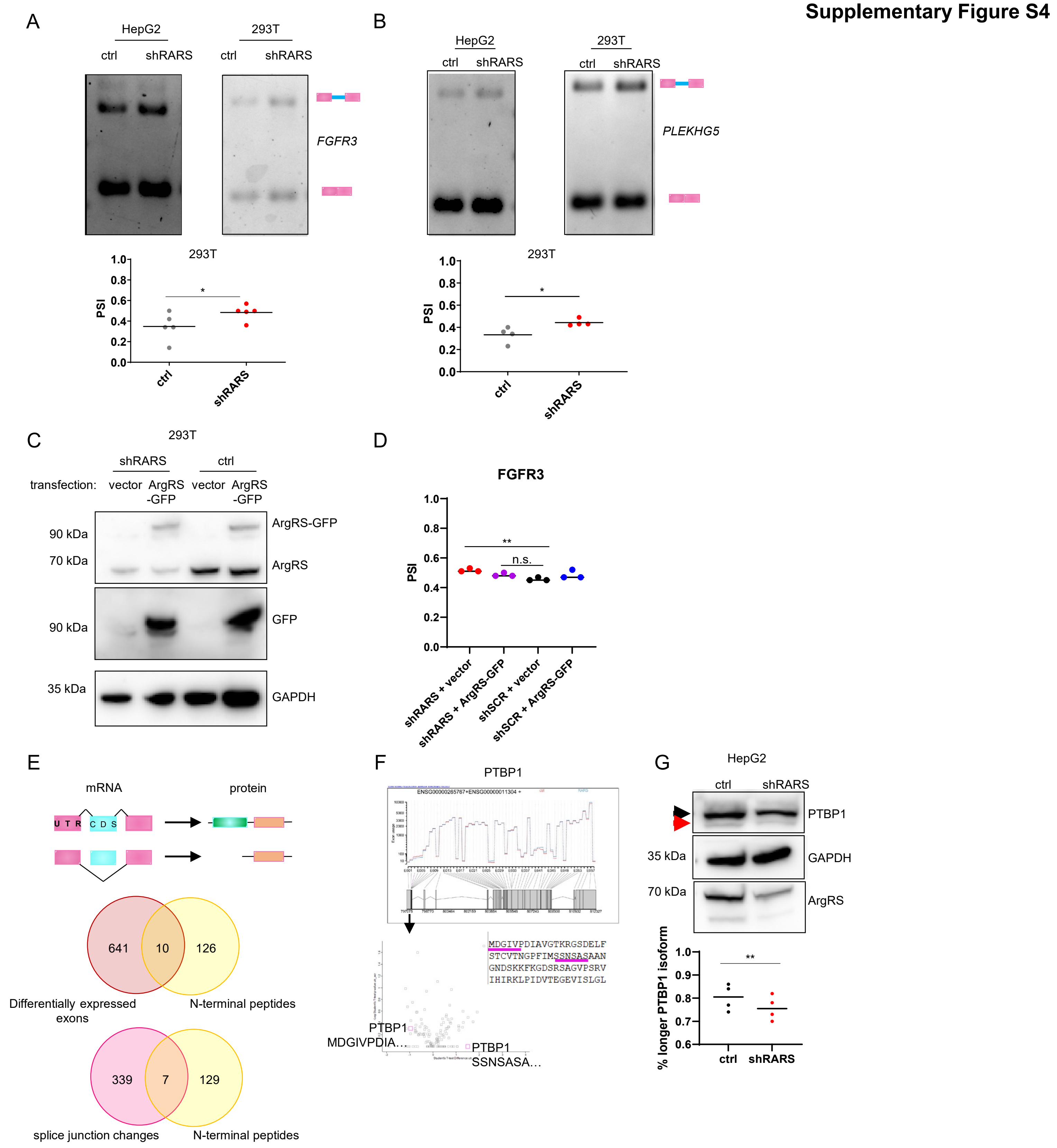


**Supplementary Figure 4: ArgRS knock down affected mRNA processing and protein isoforms.** (A, B) RT-PCR of splice isoforms for (A) *FGFR3* and (B) *PLEKHG5*. Genes were detected as alternatively spliced in HepG2 cells by RNA-seq and confirmed for HEK 293T cells. A representative image from 3 or more replicates is shown as well as the densitometric quantification for 293T. PSI: percent spliced in. Ctrl: shSCR control. p=*0.001, n=5, 5. p=*0.03, n=4, 4. (C) Rescue of ArgRS knock down by overexpression of ArgRS-GFP by transient transfection in 293T cells, shown by western blot. (D) Rescue of *FGFR3* splicing by overexpression of ArgRS-GFP. Densitometric analysis of 3 repeats is shown. p= *0.003, n=3, 3, 3, 3. (E) Genes and their corresponding proteins shared between differentially expressed exons or splice junction changes (RNA-seq) and N-terminal peptides (proteomics) in ArgRS knock down vs control cells. (F) Peptides found for PTBP1 in mass spectrometry following the enrichment of N-terminal peptides. (G) Western blot verification of preference for a higher molecular weight form of PTBP1 after ArgRS knock down. p=*0.003, n=4, 4.

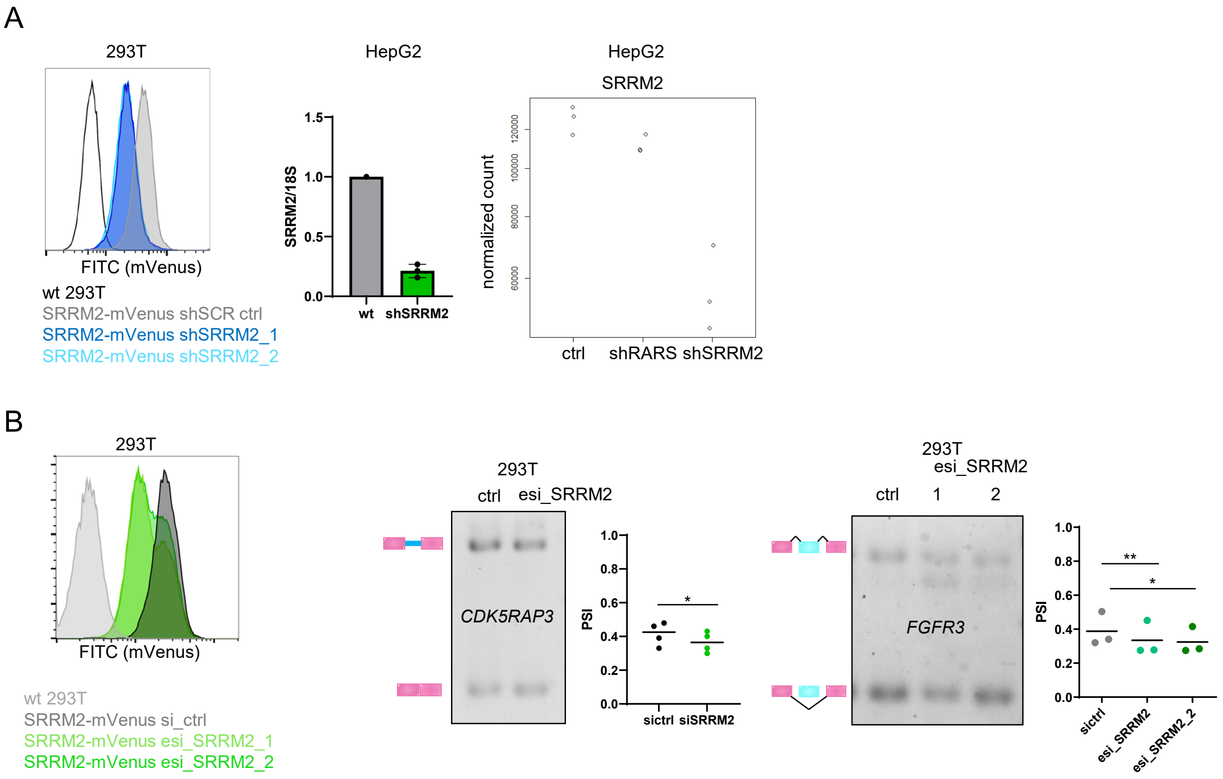


**Supplementary Figure 5: Verification of stable and transient SRRM2 knock down and its effects on mRNA splicing.** (A) Left to right: Knock down of SRRM2 in 293T_SRRM2_mVenus cells to verify shRNAs later used in HepG2 cells, assessed by flow cytometry. < 50% medium fluorescence intensity reduction upon SRRM2 knock down. Knock down of SRRM2 in HepG2 verified by qRT-PCR, three repeats, each measured in triplicates. Normalized SRRM2 counts in RNAseq. (B) Left: Verification of transient SRRM2 knock down by a heterogenous pool of siRNAs (esiRNA) in 293T_mVenus cells using flow cytometry. esi_SRRM2 1 and 2 are different batches of the same pool. ctrl: non-targeting pool of siRNAs. Right: RT-PCR splicing assays of *CDK5RAP3* and *FGFR3* upon transient knock down of SRRM2 with a pool of esiRNA in 293T cells. *CDK5RAP3*: p= *0.03, n=4, 4. *FGFR3:* p = **0.01, *0.04, n=3, 3, 3.

**Supplementary Figure S6**

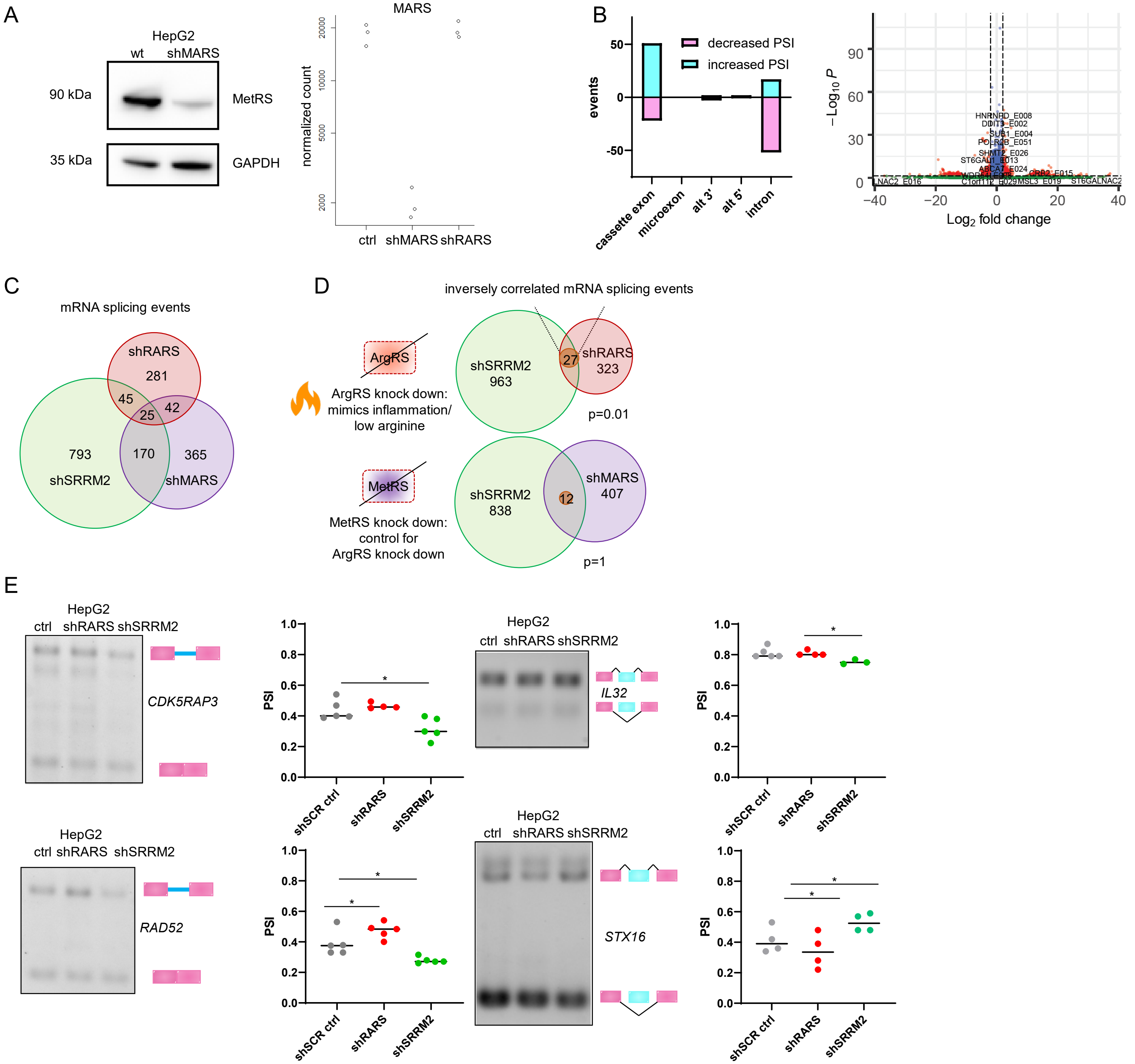


**Supplementary Figure 6: Characterization of MetRS knock down and verification of splicing events regulated by ArgRS and SRRM2 in opposite directions.** (A) Verification of MetRS knock down (shMARS) by western blot and RNAseq. (B) Splicing events with |ΔPSI| > 0.1, MV > 0 and differential exon usage (padj < 0.05). Dashed vertical lines: 2-fold change, dashed horizontal line: 1e-6. (C) Venn diagram of shared splicing event changes upon knock down of ArgRS, SRRM2, or MetRS. All splicing event changes relative to shSCR control. MV > 0, no criterion for |ΔPSI|. (D) Inversely correlated alternative splicing events shared between ArgRS and SRRM2 or MetRS and SRRM2. Hypergeometric test for overrepresentation. (E) Knock down of ArgRS and SRRM2 had opposite effects on *CDK5RAP3*, *RAD52*, and *STX16* processing as shown by RT-PCR. Representatives of 4 or 5 replicates shown with quantification by densitometry. Scheme of SRRM2 sequestration by ArgRS and resulting altered mRNA splicing. *CDK5RAP3*: p = *0.02, n= 5, 4, 5. *IL32*: p=*0.03, n=4, 4, 3. *RAD52*: p= *0.02, *0.04, n= 5, 5, 5. *STX16*: p= *0.04, *0.04, n=4, 4, 4.

### Supplementary Figure S7

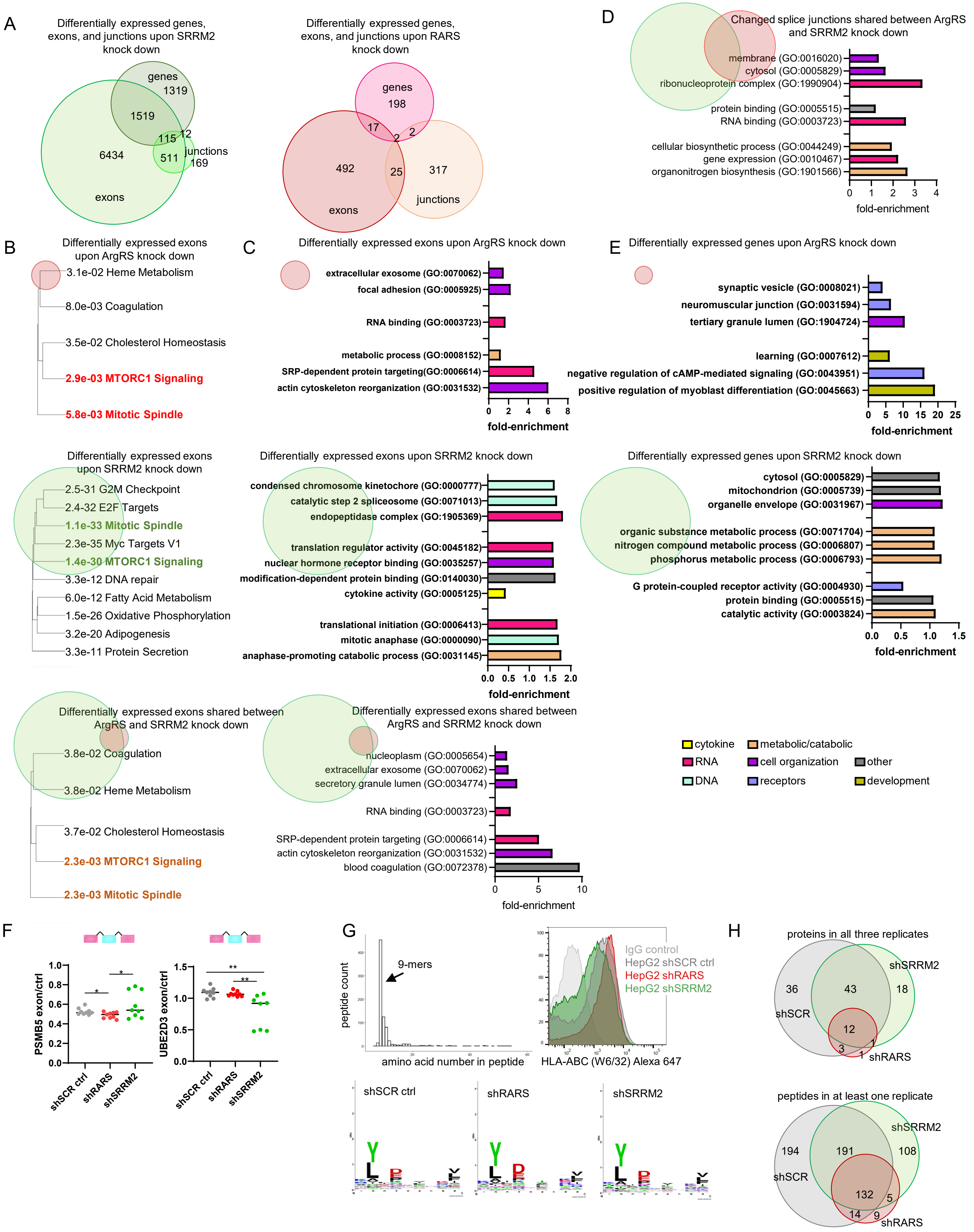

**Supplementary Figure 7: SRRM2 and ArgRS induced changes and their effects on cellular signaling.** (A) Comparison of genes with differential gene expression, differential exons, and splice junctions for shSRRM2 and shRARS. (B) Genes with differentially expressed exons enriched in MSigDB hallmark categories upon SRRM2 and ArgRS knock down (padj < 0.05) in HepG2. (C) Gene Ontology (GO) enrichment of genes with differentially expressed exons upon SRRM2 and ArgRS knock down (padj < 0.05) in HepG2. (D) GO enrichment of genes with changed splice junctions upon SRRM2 and ArgRS knock down (MV > 0) in HepG2. (C, D) All expressed genes were used as reference. (E) MSigDB hallmark categories and GO enrichment of genes with differential gene expression shared between SRRM2 and ArgRS knock down. (B-E) Circle size and Venn diagram visualize the proportion of shared exons, junctions, or genes. (F) Quantification of differentially expressed exons by qRT-PCR. PSMB5: p= *0.03, *0.04. UBE2D3: p=**0.006, **0.008. PSMB5: Splicing in ENST00000493471 led to a frameshift, resulting in PSMB5 isoform 3. UBE2D3: Skipping of ENSE00003638329 leads to an alternative start site, resulting in UBE2D3 isoform 4. (G) (top left) Length distribution histogram of MHCI peptides confirmed that isolated peptides consisted predominantly of 9 amino acid peptides. (top right) Flow cytometry of MHCI on HepG2 shSCR ctrl, ArgRS knock down, and SRRM2 knock down cells by antibody staining. (bottom). Comparable overrepresentation of amino acids at specific positions (2, 4) were found with WebLogo in 9 amino acid-long peptides. (H) Venn diagrams of identified MHCI peptides by mass spectrometry following isolation of MHCI complex, displayed as proteins corresponding to the detected peptides found in all replicates (upper panel) and peptides found in at least one replicate (lower panel).
