## Supplementary material for "Arg-tRNA synthetase links inflammatory metabolism to RNA splicing and nuclear trafficking via SRRM2": Material and Methods

Further information and requests for resources and reagents should be directed to and will be fulfilled by Prof. Paul Schimmel.

### Data and Code Availability

TGIRT-seq data are deposited in the NCBI sequence read archive and accessible through the BioProject number PRJNA561913. Poly(A) RNA-seq data are deposited at GEO (GSE165513). Mass spectrometry search results and raw data are deposited in the PRIDE archive as PXD015692 (interactomes), PXD024091 (N-terminomics), and PXD027531 (MHCI peptidome).

### Mouse model

6-8-week-old, female C57BL6/J mice from the Scripps Research breeding facility were used. Animals with the same birthdate were randomly assigned to experimental groups. All animals were sacrificed in accordance with animal protocol 19-0023 which was approved by the Scripps Research IACUC Department. Staff and researchers were wearing scrubs, facemasks, and gloves while handling animals. The rooms were kept between 69-78 degrees Fahrenheit. Individually ventilated rack caging system consisted of a cage bottom, wire bar lid, and filter top, which were sanitized in a cage wash. Feed was commercially available laboratory grade rodent diet. Purified water through reverse osmosis was provided through an automatic watering system. Cages were changed approximately every 10-14 days. Groups consisted of mice injected with (I) 100 µl 35 mg/ml murine Arginase, (II) 100 µl PBS control, (III) 50 µl 20% $CCl_4$ in olive oil, or (IV) 50 µl olive oil.

### Cell lines

HepG2 human hepatocellular carcinoma cells and human embryonic kidney (HEK) 293T cells were cultured in high glucose DMEM (Gibco) supplemented with 10% fetal bovine serum (FBS, Omega Scientific) and penicillin/streptomycin (Gibco). Both cell lines were regularly tested for mycoplasma contamination with a PCR-based, in-house assay and were consistently negative. HepG2 and 293T cells were verified by the ATCC Cell Line Authentication service using STR Profiling Results. Both cell lines were found to be a 93% match with the reference cell line profile. HepG2 cells were originally derived from a male donor and HEK 293T cells from a female embryo. Murine embryonic fibroblasts (MEF) cells were a kind gift from Yao Tong (Scripps Research). MEFs were generated from C57Bl/6J embryos around E13.5 and used at cell passage 3. The sex of the embryos that were used for MEF generation was not determined.

### Statistics

Student's t-test was used to determine significance between groups, except for mass spectrometry data for which Welch's t-test was used. Repeats are different cell passages if not noted otherwise. For densitometric analysis, paired t-test were used as repeats were run on individual gels. GraphPad Prism 8 was used for testing except for large datasets. Significance in large datasets was determined with Perseus for mass spectrometry results, DESeq2 in TGIRT-seq and RNA-seq, DEXSeq in exon analysis, and Vast-Tools in splice junction usage. Correlation coefficients were calculated with Kendall Rank correlation or Pearson correlation and enrichment of gene sets by hypergeometric testing, both in R. If not stated otherwise, significance was defined as * $p < 0.05$, ** $p < 0.01$, *** $p < 0.001$.

**Arginine depletion in mice**

To enzymatically deplete systemic arginine, mice were i.p. injected with 35 mg/kg recombinant, LPS-free (6.15 EU/mg) murine Arginase-1 purified from Clear *E. coli* (Lucigen) and sacrificed 5-7 hours later. Arginase-1 activity was tested by measuring conversion of arginine to urea. Purified Arginase-1 was mixed with a reaction buffer containing 320 mM arginine, 100 mM Hepes pH 8.5, and 10 mM $CoCl_2$. The reaction was incubated for 45 min at 37°C in a thermocycler. Urea was detected by a colorimetric assay monitoring the conversion of α-Isonitrosopropiophenone. α-Isonitrosopropiophenone was dissolved in ethanol (2 g/50 ml) and freshly diluted 1:10 in 1:1:3 water:sulphuric acid:phosphoric acid to make urea detection reagent. Urea detection reagent was mixed with the Arginase-1 reaction product or a urea standard 5:1 and incubated in a thermocycler for 60 min at 100°C followed by 15 min at 25°C.

$CCl_4$ (Sigma Aldrich) was used to elicit systemic inflammation. Mice were i.p. injected with 50 µl 20% CCl4 in olive oil, or olive oil only, and sacrificed 30 h later. In all experiments, blood was collected from the inferior vena cava with EDTA-coated syringes and plasma was separated by centrifugation at 500 x g for 3 minutes at 4°C.

**Arginine determination in plasma**

5 µl 250 µM $13C_6$-arginine (Pierce) was spiked in 20 µl plasma as an internal control. Plasma was precipitated with 200 µl 80% ice-cold methanol and dried in a speedvac. Metabolites were resuspended in 40 µl water and incubated with 20 µl freshly-made 1% Marfey's reagent in acetone (w/v) and 4 µl 1 M sodium bicarb for 1 h at 40°C in a thermocycler. Derivatized amino acids were dried in a speedvac and resuspended in 40 µl 40% methanol. Samples were measured in the Scripps Research open access mass spectrometry core using an Agilent 6125 single quadrupole mass spectrometer coupled to an Agilent 1260 LC stack. The column used was an Agilent SB-C8 5 µm 300 A 4.6x50 mm running at a flowrate of 0.5 ml/min. Mobile phases consisted of A (H2O/0.1% formic acid) and B (ACN/0.1% formic acid). Spray chamber conditions were dry gas flowrate = 12 L/min at 350 C, nebulizer pressure = 35 psi and capillary voltage = 4000 V for pos mode, and 3500 V for neg mode. 20 µl sample were injected and eluted with a 15 ml 10-35% A to B gradient. Derivatized arginine eluted at 9.3 ml. Spectra were collected in a mass range of 150-600 for both positive and negative mode and additionally in the 420-430 Da and 430-440 Da range.

**Modulation of arginine levels**

For the modulation of arginine in tissue culture, SILAC DMEM with dialyzed FBS and substituted with lysine was used as a base medium. Arginine concentrations are given as relative to 1x DMEM (0.084 g/l L-arginine monohydrochloride, dissolved in PBS). For 0x, arginine concentrations were under the detection limit of the assay described above. Cells were deprived of arginine for 4-6 hours and arginine was added back at the indicated concentrations for 2 h.

### Cell fractionation

Cultured cells were seeded at a density of 5 x $10^6$ cells per 10 cm tissue culture dish the day before (293T, MEF) or at 2.5 x $10^6$ per 10 cm tissue culture dish two days prior (HepG2) to harvest in 500 µL cell fractionation buffer (20 mM HEPES, pH 7.5, 10 mM KCl, 2 mM $MgCl_2$, 2 mM EDTA, 1 mM dithiothreitol (DTT), protease inhibitor). Spleens of arginine-depleted or control mice were homogenized by passing through a 40 µm cell sieve. Mouse liver was harvested from 4-week old C57Bl/6J mice and homogenized using a 40 µm cell sieve. The cell suspension was left on ice for 20 minutes for osmotic lysis and 100 µL were taken as whole cell lysate. The remaining cells were passed 10 times through a 27-gauge needle. Nuclei and cytoplasm were separated by centrifugation at 750x g for 5 minutes. The supernatant was taken as cytoplasmic fraction and the nuclear pellet washed three times with cell fractionation buffer. 10x RIPA buffer (0.5 M Tris/HCl, pH 7.4, 1.5 M NaCl, 2.5% Deoxycholate, 10% IGE-PAL CA-630, 10 mM EDTA) was added to 2x final concentration to all fractions. Complete lysis was achieved by freezing and thawing. Lysates were spun down at 14,000x g for 20 min to remove insoluble components and SDS loading buffer was added before western blot analysis.

### Affinity enrichment/co-immunoprecipitation

Two days prior to the experiment, 1x$10^7$ 293T or 5x$10^6$ HepG2 cells were seeded on a 15 cm tissue culture dish. Cells were lysed for 20 minutes under mild conditions in 1% IGE-PAL CA-630 (Sigma Aldrich) in Tris-buffered saline (TBS) with protease (Pierce Protein Biology) and phosphatase inhibitors (Pierce Protein Biology). Insoluble components were pelleted by centrifugation at 14,000x g for 20 minutes. Protein A/G agarose beads (30 µL, Santa Cruz Biotechnology) were equilibrated in TBS. Cell lysate was added to the beads together with 2 µL antibody against ArgRS (Biorbyt, RARS-5 (Preger et al., 2020)). For immunoprecipitation of SRRM2, antibody against SRRM2 (Santa Cruz Biotechnology) was conjugated to beads using the Co-Immunoprecipitation kit (Pierce Protein Biology) according to the manufacturer’s instructions or used as described for ArgRS (Life Technologies antibody). Immunoprecipitation was performed for 3 h (ArgRS) or overnight (SRRM2). After incubation, the beads were washed twice with TBS containing 0.1% IGE-PAL CA-630 and twice with TBS only. Precipitated proteins were either eluted using 0.1 M glycine, pH 3, for western blot or by tryptic digestion for interactome analysis.

### Western blot

Cell lysate from different cellular fractions or immunoprecipitated proteins were run on a 4-15% gradient gel (Bolt, ThermoFisher Scientific) and subsequently blotted on a PVDF membrane using an iBlot or iBlot2 device (ThermoFisher Scientific). Membranes were blocked for at least one hour with 5% bovine serum albumin (BSA) or 1% dry milk in TBS-T. Primary antibody was incubated overnight at 4°C at a 1:5000 (ArgRS: Biorbyt, GAPDH, Tubulin, and Histone H3: Cell Signaling Technology, MetRS: Abcam, GFP, PTBP1: Proteintech Group) or a 1:100 (SRRM2, Santa Cruz Biotechnology) dilution in 5% BSA/TBS-T. The next day, blots were washed three times for at least 10 minutes per wash in TBS-T and HRP-labeled secondary antibody (Invitrogen) was added at a 1:10,000 (anti-rabbit) or 1:5000 (anti-mouse) dilution in BSA/TBS-T or milk/TBS-T for at least 90 minutes. After three additional washes, membranes were developed with ProSignal Dura ECl (Genesee Scientific) and imaged with a FluorChem M (Proteinsimple).

**ArgRS interactome**

Proteomes were eluted by incubation with 25 µL 2 M urea, 5 ng/µL trypsin (Pierce Protein Biology), 1 mM DTT in 50 mM Tris/HCl, pH 7.5. After 30 minutes at room temperature, 5 mM chloroacetamide (Sigma Aldrich) was added for alkylation of cysteine residues to a total volume of 125 µL. Samples were digested overnight at 37°C and quenched with 0.5% trifluoroacetic acid (Sigma Aldrich) the next morning. Stage tip desalting was done using C18 spin tips with a 100 µL bed (Pierce Protein Biology) according to the manufacturer's instructions. The digested samples were analyzed on a Q Exactive mass spectrometer (Thermo). Approximately 250 ng of digest was injected directly onto a 20 cm, 100 µm ID column packed with Aqua 3 µm C18 resin (Phenomenex). Samples were separated at a flow rate of 400 nl/min on an Easy nLCII (Thermo). Buffer A and B were 0.1% formic acid in 5% acetonitrile and 0.1% formic acid in 80% acetonitrile, respectively. A gradient of 1-35% B over 80 minutes, an increase to 80% B over 25 minutes and held at 80% B for 5 minutes prior to returning to 1% B was used for 120 minutes total run time. Column was re-equilibrated with buffer A prior to the injection of sample. Peptides were eluted directly from the tip of the column and nanosprayed directly into the mass spectrometer by application of 2.5 kV voltage at the back of the column. The Q Exactive was operated in a data dependent mode. Full $MS^1$ scans were collected in the Orbitrap at 70 K resolution with a mass range of 400 to 1800 m/z. The 10 most abundant ions per cycle were selected for MS/MS and dynamic exclusion was used with exclusion duration of 15 seconds.

Three biological replicates were measured sequentially. Mass spectrometry data was processed with MaxQuant 1.5.7.0 (Cox and Mann, 2008) and Perseus 1.6.2.3 (Tyanova et al., 2016) as described previously (Keilhauer et al., 2015) to identify ArgRS interaction partners. In brief, raw files were searched against *Homo sapiens* reference proteome UP000005640_9606 (UniProt Consortium, 2019) (Uniprot) with the Andromeda search engine integrated in MaxQuant. Default settings were used for label-free quantification (LFQ). Phosphorylated peptides (STY) were included when searching for nuclear interaction partners of ArgRS. The resulting protein groups file was loaded into Perseus and filtered for "reverse", "potential contaminants", and "only identified by site". The log2 values of LFQ intensities were calculated and all proteins with less than two valid values/group discarded. Missing values were replaced from a normal distribution (width 0.3, downshift 1.8) and Welch's t-test was used to calculate t-test significance and difference.

**Generation of SRRM2-mVenus cell line**

A gRNA (CCAUGAGACACCGCUCCUCC) targeted to the second last exon of SRRM2 (exon 14) was inserted into pSpCas9(BB)-2A-GFP. pSpCas9(BB)-2A-GFP (PX458) was a gift from Feng Zhang, Addgene plasmid #48138; http://n2t.net/addgene:48138; RRID:Addgene_48138 (Ran et al., 2013)). mVenus (mVenus C1) was a gift from Steven Vogel, Addgene plasmid #27794; http://n2t.net/addgene:27794; RRID:Addgene_27794 (Koushik et al., 2006). The donor vector for insertion of mVenus was based on pcDNA6.

The CMV promoter and multiple cloning site of pcDNA6 were substituted for mVenus flanked by 800 bp upstream and downstream of the Cas9 cleavage site. Cells were transfected with both plasmids at a 1:2 ratio gRNA/Cas9:donor vector and sorted for GFP expression after two days (successful expression of pSpCas9-GFP) by the Flow Cytometry core at Scripps Research. Single cells were expanded into monoclonal cell lines and evaluated for mVenus expression after expansion by flow cytometry. About 1/3 of all tested clones showed mVenus fluorescence. Correct genomic integration was assessed by PCR on isolated genomic DNA (DNeasy Blood and Tissue Kit, Qiagen, Supplementary Figure S2J). PCR primers: GTGGTGCCTGAGGTGGTGCC/CCACTCCCAAAATGGGGCCG

Successful labeling of SRRM2 only was confirmed by the reduction of mVenus by two individual shRNAs directed against SRRM2 (Supplementary Figure S5A).

**ArgRS, MetRS, and SRRM2 knock down**

shRNAs against ArgRS (shRARS_1: gtggacacaagcataagtaaa, shRARS_2: ggagcagttacaagaagaaaa) were cloned into a pLKO.1 vector (pLKO.1 - TRC cloning vector was a gift from David Root, Addgene plasmid #10878; http://n2t.net/addgene:10878; RRID:Addgene_10878 (Moffat et al., 2006)). A non-targeting control shRNA pLKO.1 shSCR was a gift from Sheila Stewart (Addgene plasmid #17920; http://n2t.net/addgene:17920; RRID:Addgene_17920 (Saharia et al., 2008)). shRNAs against *SRRM2* and *MARS* were acquired from The RNAi Consortium *via* Sigma Aldrich (shSRRM2_1: cgccacctaaacagaaatc, shSRRM2_2: gttgggactggaggttgta, shMARS_1: caaggaaacattgtccgagaac, shMARS_2: tcgacatggcaaccaatatatc). Plasmids were co-transfected into 293T cells together with lentiviral packaging plasmids pRSV-rev, pMDLg/pRRE, and pMD2.G (plasmids were a gift from Didier Trono (Addgene plasmid #12253 and #12251; http://n2t.net/addgene:12253 and :12251; RRID:Addgene_12253 and _12251 (Dull et al., 1998)), and a gift from the Torbett lab at Scripps Research) using Lipofectamine 2000 (Invitrogen). Medium was changed after overnight transfection and collected two days later. Supernatant containing viral particles was filtered through a 0.45 µm syringe filter, and either used directly or stored at -80°C. For viral transduction, SRRM2-mVenus-tagged 293T or HepG2 cells were seeded a day prior on 6 cm tissue culture dishes and incubated with 1 ml undiluted viral particles in the presence of 8 µg polybrene (EMD Millipore) for two hours. Afterwards, fresh medium was added, and the infection left to proceed for two days. Successfully transduced cells were selected with 10 µg/ml (293T) or 50 µg/ml (HepG2) puromycin (AdipoGen Life Science). ArgRS knock down was confirmed via western blot and quantitative real-time PCR.

**Reverse transcription and quantitative real-time PCR (RT-qPCR)**

RNA was isolated using Trizol (Invitrogen) according to the manufacturer's instructions. 500 ng RNA was reverse-transcribed using MultiScribe Reverse Transcriptase (Applied Biosystems) according to instructions in the High-Capacity cDNA Reverse Transcription Kit (Applied Biosystems) with random hexamers (Qiagen) as primers. qRT-PCR was performed with Power SYBR Green Master Mix (Thermo Fisher Scientific) on a StepOnePlus (Applied Biosystems). The following primers were used:
*RARS*: GAGAGTATAAGCCGCCTCTTTG/TCCTCCTCAGTATCAAACCTCT;
*SRRM2*: TGACAGCAAATCTCGACTATCC/GGTTCAGGAGAGGAATCAGAAC

18S: CTGAGAAACGGCTACCACATC/GCCTCGAAAGAGTCCTGTATTG

**Immunofluorescence**

Glass coverslips (12 mm, 1.5 thickness, Neuvitro Corporation) were sterilized with 70% ethanol, coated with 0.01% Collagen I (MP Biomedicals Inc) overnight, and stored in PBS until use. Two days prior to staining, 10,000 HepG2 or 293T cells were seeded on coverslips. Cells were washed, fixed with 4% PFA in PBS for 20 minutes, permeabilized in 0.25% Triton X-100 for 5 minutes, blocked with 1% FCS, 2% BSA in PBS (IF buffer) for a minimum of 15 minutes, and stained overnight with anti-SRRM2 antibody or anti-ArgRS antibody (both SCBT, 1:100, all dilutions in IF buffer) at 4°C. Cells were washed twice with PBS and once with IF buffer between all steps. Anti-mouse AlexaFluor 488 secondary antibody (Abcam) was diluted 1:250 and added to the cells for 45 minutes at room temperature. Anti-SC35/SFRS2 (Proteintech), Methionine-tRNA Synthetase (MetRS, Abcam), or Arginyl-tRNA Synthetase antibody (Biorbyt) were added at a 1:100 dilution (ArgRS) or 1:200 dilution (MetRS, SC35) and incubated at room temperature for 3 hours. Cross-adsorbed anti-rabbit Alexa Fluor 568 secondary antibody (Abcam) was added for 45 minutes at room temperature at a dilution of 1:250. Hoechst 33342 (AdipoGen Life Science) was added at 1:500 in PBS for 15 minutes. Cover slips were washed in PBS, dipped in water, mounted on soft-set Vectashield (Vectorlabs), and sealed with gel glue or nailpolish. Slides were stored at 4°C until imaging. Staining of SRRM2 with a monoclonal antibody raised against a C-terminal peptide of SRRM2 showed distinct nuclear structures, strongly reminiscent of an SRRM2 antibody staining deposited in the Human Protein Atlas with a different antibody (HPA066181).

**Confocal microscopy and colocalization**

Microscopes were maintained by the Core Microscopy Facility at Scripps Research. For co-localization of SRRM2, ArgRS, SFPQ, and MetRS, a Zeiss LSM 880 Airyscan confocal laser scanning microscope with a 100x oil objective was used for imaging in airyscan mode. Confocal images were recorded with a pixel dwell time of 1.5 µs in 16-bit depth with a scaling of 0.034 µm (x, y) and 0.145 µm (z). Z-stacks covering the nucleus were taken with a motorized stage. Images were processed with “Airyscan Processing” in Zen (Zeiss). Estimation of colocalization was done with Imaris (Oxford Instruments). First, the nuclear volume of each cell was determined by using Hoechst DNA staining as a mask to delineate the cell nucleus, with an adequate threshold to filter spillover of Hoechst signal. Second, the nuclear signal of two antibody stains were colocalized by comparing nuclear volumes in individual cells and the percentage of colocalized volume was noted as well as Manders overlap coefficients and Pearson correlation coefficient. Manders overlap coefficient calculates the overlap between two colors from their absolute intensities and results in individual coefficients for each color (co-occurrence of color1 with color2 and vice versa). In contrast, Pearson correlation coefficient uses the deviation from the mean and results therefore in one coefficient for the correlation between both colors (relationship between the signal intensities). Background for the antibody staining was determined by comparison of signal intensity to a control in which cells were stained with secondary antibody only or by Costes thresholding in Imaris, if stated so in the figure legend. Videos demonstrating colocalization (Supplementary Videos 1-4) were recorded with Imaris (Oxford Instruments). Supplementary Figure S2K was taken on a Zeiss LSM

710 confocal laser scanning microscope with a 63x oil objective. Cells were fixed in 4% PFA/PBS for 10 minutes to retain mVenus fluorescence.

**Fluorescence recovery after photobleaching (FRAP)**

SRRM2-mVenus 293T cells expressing either a non-targeting control shRNA (shSCR ctrl) or shRNA against ArgRS (shRARS) were seeded on collagen-I coated (as described above for cover slips) glass bottom 35 mm dishes with 1.5 thickness (World Precision Instrument) two days prior to imaging. Live cell imaging was performed on a Zeiss LSM 780 or a LSM 880 confocal laser scanning microscope at 37°C, 5% $CO_2$ with a 63x water objective. Fluorescent bleaching was achieved by 20 cycles of 100% intensity at 514 nm. Pictures were taken every 2 s for 75 cycles with a pixel dwell time of 1.58 µs and fluorescence in regions of interest (ROIs) were recorded with ZEN (Zeiss). The resulting data was subtracted from background, divided by fluorescence intensity of the same area before bleaching (normalization), and the geometric mean of ROIs was fitted with a one-exponential model to calculate the half-time of fluorescence recovery (GraphPad Prism 5). The mean of individual measurements was calculated and plotted together with standard error means. Pictures were exported with Fiji/ImageJ. Immobile fractions were calculated from dedicated measurements with images taken every 10 s (as opposed to every 2 s for the calculation of half-time of recovery) to minimalize photobleaching during imaging (at the cost of temporal resolution). FRAP recordings were excluded if sample drift was noted during the recording. Significance was tested with a two-tailed Student's t-test (GraphPad Prism 5). We compared SRRM2-mVenus fluorescence recovery half-times to known literature values of ASF/SRSF1 (Phair and Misteli, 2000) recovery. Secondary structure prediction of both proteins was carried out with Sable (Adamczak et al., 2004).

**Flow cytometry and fluorescence activated cell sorting**

Flow cytometry was performed on instruments maintained by the Flow Cytometry core at Scripps Research. 293T cells were detached using Trypsin/EDTA and resuspended in DMEM with 10% FBS. The cell suspension was mixed 1:1 with sorting buffer (2.5 mM EDTA, 25 mM HEPES, pH 7.0, 1% FBS, 1% penicillin/streptomycin in PBS) and analyzed on an LSR II analytical flow cytometer (BD Bioscience). mVenus fluorescence was detected in the FITC channel (filter 525/50) after excitation by a 488 nm laser. Flow cytometry results were analyzed with FlowJo version 10.06 or higher. Fluorescence activated cell sorting was performed by members of the Scripps Research Flow Cytometry core on a MoFlo Astrios EQ jet-in-air sorting flow cytometer (Beckman Coulter). Two days after transfection, cells were detached using Accutase (Stemcell Technologies), resuspended in DMEM with FBS, and spun down to remove serum to prevent cells from adhering. Following a wash with PBS, cells were diluted in sorting buffer to $1x10^6$ cells/ml. Single cells were sorted into a 96 well plate prefilled with 100 µL medium and expanded to monoclonal cell lines. For quantification of MHCI expression, cells were detached with EDTA and stained with 1 ug/ml HB-95 W6/32 for 20 minutes on ice followed by Alexa 647-cojugated anti-mouse secondary antibody for 20 minutes on ice, both in sorting buffer.

**Cell viability**

Cell viability and proliferation was assessed by measuring AlamarBlue (AbD Serotec) conversion. HepG2 or 293T cells were seeded in 96-well plates (10,000 cells/well) and 1:100 AlamarBlue was added two hours prior to measurement on a Synergy H1 (Biotek, $\lambda_{exc}$ 530 nm, $\lambda_{em}$ 580 nm). All measurements were done in technical quadruplicates.

### RNA isolation and TGIRT-seq library preparation

$2.5x10^6$ HepG2 cells expressing either a non-targeting shRNA control (shSCR) or a shRNA directed against ArgRS (gene name: *RARS*, shRARS) were seeded on 10 cm tissue culture dishes two days prior to RNA isolation. Cells were starved off arginine for 6 hours and supplemented with 10x arginine two hours prior to harvest. RNA was isolated by using a mirVana miRNA isolation kit (Invitrogen) with phenol extraction following the manufacturer's instruction for total RNA isolation. RNA integrity was evaluated with a Bioanalyzer (Agilent). Extracted total RNA was treated with TURBO DNase (Thermo Fisher) according to the manufacturer's instruction, and the DNase-treated RNA was then ribo-depleted by using a Ribo-Zero Gold (Human/Mouse/Rat) kit (Illumina). For each library, 50 ng of ribo-depleted RNA was fragmented by using an RNA Fragmentation Module kit (New England Biolabs) to a median size of 55-75 nt, and 3' phosphates were removed by treatment with T4 polynucleotide kinase (Epicentre).

TGIRT-seq libraries were prepared essentially as described (Qin et al., 2016) using a modified R2R adapter that decreases adapter dimer formation (Xu et al., 2019). Reverse transcription reactions contained fragmented, dephosphorylated cellular RNA, 100 nM R2 RNA/R2R DNA starter duplex, and 1 µM TGIRT-III (InGex) in reaction buffer (20 mM Tris-HCl, pH 7.5, 450 mM NaCl, 5 mM $MgCl_2$, 5 mM DTT). Reactions were pre-incubated at room temperature for 30 min and then initiated by addition of 1 mM dNTPs (an equimolar mix of 1 mM dATP, dCTP, dGTP, and dTTP) and raising the temperature to 60°C. Reverse transcription by TGIRT-III is initiated by template switching from a starting duplex consisting of a 35 nt DNA primer encoding the reverse complement of the Illumina Read 2 sequencing primer binding site (R2R) annealed to a 34-nt complementary RNA oligonucleotide (R2), leaving a single nucleotide 3' DNA overhang composed of an equimolar mixture of A, G, C and T. The RNA oligonucleotide is blocked at its 3' end with C3Sp (IDT) to inhibit template switching to itself. Reactions were incubated at 60°C for 15 min and terminated by adding 5 N NaOH to a final concentration of 0.25 N and incubating at 95°C for 3 min to degrade RNAs and denature protein. The reactions were then cooled to room temperature and neutralized with 5 N HCl. cDNAs were purified by using a Qiagen MinElute Reaction Cleanup Kit and then ligated at their 3' ends to a DNA oligonucleotide encoding the reverse complement of the Illumina Read 1 primer binding site (R1R) using Thermostable 5' AppDNA/RNA Ligase (New England Biolabs). Ligated cDNAs were re-purified with a MinElute Reaction Cleanup Kit and amplified by PCR for 12 cycles using Phusion DNA polymerase (Thermo Fisher Scientific) with overlapping multiplex and index primers that add sequences necessary for Illumina sequencing. PCR products were purified with AMPure XP beads (Beckman-Coulter) to remove unused PCR primers and adaptor dimers. Libraries were sequenced on a NextSeq 500 instrument (2x75 nt, paired end reads) at the Genomic Sequencing and Analysis Facility at the University of Texas at Austin.

### TGIRT-seq bioinformatic analysis

After sequencing, Fastq files were processed by using the TGIRT-map pipeline (Wu et al., 2018). Briefly, the sequencing adapters were removed with Atropos (Didion et al., 2017) using options *trim -U 1 --minimum-length=15 --threads=24 --no-cache-adapters --error-rate=0.1 –b AAGATCGGAAGAGCACACGTCTGAACTCCAGTCAC -B GATCGTCGGACTGTAGAACTCTGAACGTGTAGA.* Trimmed reads were then aligned against 5S rRNA (GenBank accession: X12811.1), complete rRNA repeat unit (GenBank accession U13369.1) and hg19 Genomic tRNA database (Chan and Lowe, 2016) using BOWTIE2 (Langmead and Salzberg, 2012). The remaining unaligned reads were then sequentially aligned to the human genome (hg19) using a splice-aware aligner (HISAT2) (Kim et al., 2015) and a sensitive local aligner (BOWTIE2) (Langmead and Salzberg, 2012). Finally, gene quantification was done on genomic loci, except for tRNA and rRNA, which required an additional step of re-aligning and counting (Supplementary Table S14). Differential gene expression analysis was done using DESeq2 (Love et al., 2014). Exon quantification was done by FeatureCount (Liao et al., 2014) and differential exon usage was quantified using DEXSeq (Anders et al., 2012). DEXSeq uses a negative binomial model to test for differential exon usage between two samples while normalizing for the differences in gene expression (Anders et al., 2012). Gene names were assigned using biomaRt (Durinck et al., 2009).

**Poly(A) RNA-seq library preparation and bioinformatic analysis**

Poly(A) RNA-seq libraries were prepared and sequenced by the Genomic Sequencing and Analysis Facility at the University of Texas at Austin. RNAs were selected by using a Poly(A) Purist MAG Kit (Thermo Fisher, AM1922) using the manufacturer's protocol followed by library preparation using a NEBNext Ultra II Directional RNA Library Kit for Illumina (New England Biolabs, E7760), also following the manufacturer's protocol. Samples were sequenced on a NovaSeq 6000 instrument using SP flowcells (2x150 nt, paired end reads). Reads were trimmed with trimgalore and aligned to grch37 with STAR (Dobin et al., 2013) (Supplementary Table S15). Counts were generated with Subread (Liao et al., 2014). Differential exon usage analysis was performed with DEXSeq (Anders et al., 2012). If exons were assigned to overlapping genes, gene symbols were assigned to the first gene and counted as one gene except in the direct comparison between DEXSeq results and N-terminal peptides, were all genes were used. Differentially expressed genes were identified with DESeq2 (Love et al., 2014). Enrichment analysis was performed using Panther (Mi et al., 2019) and ShinyGO (Ge et al., 2020). For Panther, all genes mapped to grch37 were used as a reference list. Vast-tools was used to identify differentially spliced events, including cassette exons, microexons, 3' and 5' alternative splice sites, and retained splice sites (Irimia et al., 2014). Untrimmed reads were first aligned to the hg19 assembly and PSI values for all annotated splicing events were measured for each sample. Events were considered if, in at least of half of the profiled samples, they had ≥ 15 reads overlapping the included/excluded splice junctions and a ratio of reads mapping to the upstream and downstream junctions (a balance score) of < 2 or 2 to 5. These events were than processed through the vast-tools diff module (Han et al., 2017), which calculates the predicted ΔPSI between samples and assigns a minimum ΔPSI value (MV), with a ≥ 0.95 probability, for each event. For example, MV > 0 |ΔPSI| ≥ 0.1 indicates ≥ 0.95 probability of at least a 10% difference in splice junction usage in either direction.

### RT-PCR for verification of splicing events

RNA was reverse transcribed and amplified using OneStep RT-PCR (Qiagen) according to the manufacturer's instructions but in a reduced volume (10 µl final). 40 ng of HepG2 RNA or 100 ng of 293T RNA were transcribed per reaction. Primers were designed using VastDB (Tapial et al., 2017) for splice junction changes and manually for differential exon usage. Amplified regions of different splice isoforms were separated on a 2.5% agarose/TAE gel and visualized after staining with SybrSafe on a Biorad ChemiDoc Imager.

### Detection of N-terminal peptides

A modified protocol for the Terminal amine isotopic labeling of substrate (TAILS) was used to enrich for N-terminal peptides (Kleifeld et al., 2011). Cells were lysed, proteins were extracted by a methanol/chloroform precipitation, and N-termini were blocked overnight with formaldehyde. Proteins were extracted again with methanol/chloroform to avoid carry-over of reactive formaldehyde and digested with trypsin overnight. HPG-ALD Polymers were acquired through Flintbox and added to the peptides at a 1:50 ratio to deplete peptides with unblocked reactive N-termini derived from the trypsin digest. The next day, the reaction was quenched, and free peptides separated from the polymer with a 10 kDa spin filter. Peptides were fractionated with Pierce High pH Reversed-Phase Peptide Fractionation kit (Thermo) according to the manufacturer's instructions, all 9 fractions were dried down, resuspended in 5% acetonitrile and 0.1% formic acid and were analyzed on the Q Exactive mass spectrometer as described above. Raw data was processed with MaxQuant (Cox and Mann, 2008) with dimethyllysine and dimethylated N-termini as a fixed modification. Peptides were searched against a reference proteome (Uniprot) with free N-termini and ArgC as protease and label free quantification was used with default settings. The resulting intensities and LFQ values were filtered in Perseus (Tyanova et al., 2016) against a contaminant database. Reverse and peptides preceded by arginine, which are likely derived from trypsin cleavage, were excluded. MaxQuant (version 1.6.7.0) was run on the Scripps High Performance Computing core. Genes with overlapping differential exon usage (DEXSeq padj < 0.05) and N-terminal peptides (Student's t-test difference > $|\Delta 1|$) were further manually analyzed to identify the position of the differential exon in the gene and the resulting peptide sequence.

### IDH and proteasome activity

10,000 HepG2 cells were seeded in white 96 well plates two days prior to both assays. For IDH activity, cells were lysed in co-immunoprecipitation lysis buffer and mixed with an end concentration of 100 µM NADP, 2 mM $MnCl_2$, and 5 mM isocitrate. IDH activity was measured by monitoring fluorescence at 460 nm after excitation at 340 nm, once per minute for 10 minutes at 37°C. Proteasomal subunit β5 activity was measured using the ProteasomeGlo assay (Promega) according to the manufacturer's instructions in the same format. Total protein concentration (IDH, with BCA assay) or cell count (β5 activity, with alamar blue assay) were used to normalize between treatment conditions.

### MHCI peptidome

$5x10^8$ cells were seeded per replicate 2 days prior to harvest and frozen at -80°C until MHCI peptide enrichment. HB-95 W6/32 hybridoma were acquired by ATCC, cultured in DMEM with 10% FBS, and diluted in HyClone CDM4MAb serum free media stepwise till 10 % DMEM was reached. Cells were left to condition the serum-reduced media for 5 days. 300 ml of medium were loaded on 3 ml of Protein A, washed with PBS, and eluted with 100 mM glycine, pH 3.0, into a final concentration of 130 mM Hepes, pH 7.5 in 5 column volumes. Coupling of antibody to protein A was performed as described previously (Purcell et al., 2019). Cell lysis followed instructions by Purcell et al., but centrifugation speed was changed to 40,000 x g for 30 minutes. Separation of MHCI from MHCI-bound peptides was performed as described previously (Bassani-Sternberg et al., 2015). Peptides were measured on a QE-HFX and searched against the human proteome with ProLuCID (Xu et al., 2015) without protease specificity and no requirements for tryptic peptides for protein identification. The percentage of arginine in the proteins from which the MHCI peptides originated was not significantly changed between groups. MHCI immunogenicity score was calculated as described previously (Calis et al., 2013).

**References:**


Adamczak, R., Porollo, A., and Meller, J. (2004). Accurate prediction of solvent accessibility using neural networks-based regression. Proteins *56*, 753–767.

Anders, S., Reyes, A., and Huber, W. (2012). Detecting differential usage of exons from RNA-seq data. Genome Res. *22*, 2008–2017.

Bassani-Sternberg, M., Pletscher-Frankild, S., Jensen, L.J., and Mann, M. (2015). Mass spectrometry of human leukocyte antigen class I peptidomes reveals strong effects of protein abundance and turnover on antigen presentation. Mol. Cell. Proteomics MCP *14*, 658–673.

Calis, J.J.A., Maybeno, M., Greenbaum, J.A., Weiskopf, D., De Silva, A.D., Sette, A., Keşmir, C., and Peters, B. (2013). Properties of MHC class I presented peptides that enhance immunogenicity. PLoS Comput. Biol. *9*, e1003266.

Chan, P.P., and Lowe, T.M. (2016). GtRNAdb 2.0: an expanded database of transfer RNA genes identified in complete and draft genomes. Nucleic Acids Res. *44*, D184-189.

Cox, J., and Mann, M. (2008). MaxQuant enables high peptide identification rates, individualized p.p.b.-range mass accuracies and proteome-wide protein quantification. Nat. Biotechnol. *26*, 1367–1372.

Didion, J.P., Martin, M., and Collins, F.S. (2017). Atropos: specific, sensitive, and speedy trimming of sequencing reads. PeerJ *5*, e3720.

Dobin, A., Davis, C.A., Schlesinger, F., Drenkow, J., Zaleski, C., Jha, S., Batut, P., Chaisson, M., and Gingeras, T.R. (2013). STAR: ultrafast universal RNA-seq aligner. Bioinforma. Oxf. Engl. *29*, 15–21.

Dull, T., Zufferey, R., Kelly, M., Mandel, R.J., Nguyen, M., Trono, D., and Naldini, L. (1998). A third-generation lentivirus vector with a conditional packaging system. J. Virol. *72*, 8463–8471.

Durinck, S., Spellman, P.T., Birney, E., and Huber, W. (2009). Mapping identifiers for the integration of genomic datasets with the R/Bioconductor package biomaRt. Nat. Protoc. *4*, 1184–1191.

Ge, S.X., Jung, D., and Yao, R. (2020). ShinyGO: a graphical gene-set enrichment tool for animals and plants. Bioinforma. Oxf. Engl. *36*, 2628–2629.

Han, H., Braunschweig, U., Gonatopoulos-Pournatzis, T., Weatheritt, R.J., Hirsch, C.L., Ha, K.C.H., Radovani, E., Nabeel-Shah, S., Sterne-Weiler, T., Wang, J., et al. (2017). Multilayered Control of Alternative Splicing Regulatory Networks by Transcription Factors. Mol. Cell *65*, 539-553.e7.

Irimia, M., Weatheritt, R.J., Ellis, J.D., Parikshak, N.N., Gonatopoulos-Pournatzis, T., Babor, M., Quesnel-Vallières, M., Tapial, J., Raj, B., O'Hanlon, D., et al. (2014). A highly conserved program of neuronal microexons is misregulated in autistic brains. Cell *159*, 1511–1523.

Keilhauer, E.C., Hein, M.Y., and Mann, M. (2015). Accurate protein complex retrieval by affinity enrichment mass spectrometry (AE-MS) rather than affinity purification mass spectrometry (AP-MS). Mol. Cell. Proteomics MCP *14*, 120–135.

Kim, D., Langmead, B., and Salzberg, S.L. (2015). HISAT: a fast spliced aligner with low memory requirements. Nat. Methods *12*, 357–360.

Kleifeld, O., Doucet, A., Prudova, A., auf dem Keller, U., Gioia, M., Kizhakkedathu, J.N., and Overall, C.M. (2011). Identifying and quantifying proteolytic events and the natural N terminome by terminal amine isotopic labeling of substrates. Nat. Protoc. *6*, 1578–1611.

Koushik, S.V., Chen, H., Thaler, C., Puhl, H.L., and Vogel, S.S. (2006). Cerulean, Venus, and VenusY67C FRET reference standards. Biophys. J. *91*, L99–L101.

Langmead, B., and Salzberg, S.L. (2012). Fast gapped-read alignment with Bowtie 2. Nat. Methods *9*, 357–359.

Liao, Y., Smyth, G.K., and Shi, W. (2014). featureCounts: an efficient general purpose program for assigning sequence reads to genomic features. Bioinforma. Oxf. Engl. *30*, 923–930.

Love, M.I., Huber, W., and Anders, S. (2014). Moderated estimation of fold change and dispersion for RNA-seq data with DESeq2. Genome Biol. *15*, 550.

Mi, H., Muruganujan, A., Ebert, D., Huang, X., and Thomas, P.D. (2019). PANTHER version 14: more genomes, a new PANTHER GO-slim and improvements in enrichment analysis tools. Nucleic Acids Res. *47*, D419–D426.

Moffat, J., Grueneberg, D.A., Yang, X., Kim, S.Y., Kloepfer, A.M., Hinkle, G., Piqani, B., Eisenhaure, T.M., Luo, B., Grenier, J.K., et al. (2006). A lentiviral RNAi library for human and mouse genes applied to an arrayed viral high-content screen. Cell *124*, 1283–1298.

Phair, R.D., and Misteli, T. (2000). High mobility of proteins in the mammalian cell nucleus. Nature *404*, 604–609.

Preger, C., Wigren, E., Ossipova, E., Marks, C., Lengqvist, J., Hofström, C., Andersson, O., Jakobsson, P.-J., Gräslund, S., and Persson, H. (2020). Generation and validation of recombinant antibodies to study human aminoacyl-tRNA synthetases. J. Biol. Chem. *295*, 13981–13993.

Purcell, A.W., Ramarathinam, S.H., and Ternette, N. (2019). Mass spectrometry-based identification of MHC-bound peptides for immunopeptidomics. Nat. Protoc. *14*, 1687–1707.

Qin, Y., Yao, J., Wu, D.C., Nottingham, R.M., Mohr, S., Hunicke-Smith, S., and Lambowitz, A.M. (2016). High-throughput sequencing of human plasma RNA by using thermostable group II intron reverse transcriptases. RNA N. Y. N *22*, 111–128.

Ran, F.A., Hsu, P.D., Wright, J., Agarwala, V., Scott, D.A., and Zhang, F. (2013). Genome engineering using the CRISPR-Cas9 system. Nat. Protoc. *8*, 2281–2308.

Saharia, A., Guittat, L., Crocker, S., Lim, A., Steffen, M., Kulkarni, S., and Stewart, S.A. (2008). Flap endonuclease 1 contributes to telomere stability. Curr. Biol. CB *18*, 496–500.

Tapial, J., Ha, K.C.H., Sterne-Weiler, T., Gohr, A., Braunschweig, U., Hermoso-Pulido, A., Quesnel-Vallières, M., Permanyer, J., Sodaei, R., Marquez, Y., et al. (2017). An atlas of alternative splicing profiles and functional associations reveals new regulatory programs and genes that simultaneously express multiple major isoforms. Genome Res. *27*, 1759–1768.

Tyanova, S., Temu, T., Sinitcyn, P., Carlson, A., Hein, M.Y., Geiger, T., Mann, M., and Cox, J. (2016). The Perseus computational platform for comprehensive analysis of (prote)omics data. Nat. Methods *13*, 731–740.

UniProt Consortium (2019). UniProt: a worldwide hub of protein knowledge. Nucleic Acids Res. *47*, D506–D515.

Wu, D.C., Yao, J., Ho, K.S., Lambowitz, A.M., and Wilke, C.O. (2018). Limitations of alignment-free tools in total RNA-seq quantification. BMC Genomics *19*, 510.

Xu, H., Yao, J., Wu, D.C., and Lambowitz, A.M. (2019). Improved TGIRT-seq methods for comprehensive transcriptome profiling with decreased adapter dimer formation and bias correction. Sci. Rep. *9*, 7953.

Xu, T., Park, S.K., Venable, J.D., Wohlschlegel, J.A., Diedrich, J.K., Cociorva, D., Lu, B., Liao, L., Hewel, J., Han, X., et al. (2015). ProLuCID: An improved SEQUEST-like algorithm with enhanced sensitivity and specificity. J. Proteomics *129*, 16–24.

**Supplementary Figures 1-7**

**Supplementary tables**

**Table S1**: ArgRS nuclear interactome in HepG2.

**Table S2**: ArgRS interactome in HepG2 whole cell lysate.

**Table S3**: ArgRS interactome in 293T whole cell lysate.

**Table S4**: MetRS interactome in 293T whole cell lysate.

**Table S5**: Proteins shared between interactomes.

**Table S6**: tRNA expression upon ArgRS knock down as calculated from TGIRTseq.

**Table S7**: Splice junctions with MV > 0 and |ΔPSI| > 0.1 upon ArgRS knock down.

**Table S8**: Differentially expressed exons upon ArgRS knock down (padj > 0.05).

**Table S9**: Peptides identified after enrichment of N-terminal peptides with |pDiff| > 1 upon ArgRS knock down.

**Table S10**: Splice junctions with MV > 0 and |ΔPSI| > 0.1 upon SRRM2 knock down.

**Table S11**: Differentially expressed exons upon SRRM2 knock down (padj > 0.05).

**Table S12**: Splice junctions with MV > 0 and |ΔPSI| > 0.1 upon MetRS knock down.

**Table S13**: Genes associated with Hallmark and GO terms that are inversely modulated by ArgRS and SRRM2.

**Table S14**: Mapping statistics for TGIRT-seq. Numbers on the top of each cell represent numbers of read pairs. Numbers within parentheses represent percentages. Raw pairs indicate the number of read pairs for the sample after Illumina sample demultiplexing. Trimmed pairs indicate the number of read pairs that passed adapter- and quality-trimmings, and length-cutoffs, number in parentheses indicates percentage of trimmed pairs respect to raw pairs. tRNA/rRNA pairs indicates number of read pairs that can be concordantly aligned to ribosomal RNA repeat units (GenBank accession numbers: X12811.1 and U13369.1)) or tRNA (GtRNAdb; (Chan and Lowe, 2016), percentage is computed respect to trimmed pairs. Non tRNA/rRNA pairs indicate the number of read pairs that are not mappable to rRNA or tRNA, percentage is computed respect to trimmed pairs. HISAT2 mapped pairs indicates number of read pairs that can be aligned concordantly to human genome (hg19) by HISAT2 (Kim et al., 2015), percentage is computed respect to non tRNA/rRNA pairs. BOWTIE2 mapped pairs indicates number of read pairs that aligned to human genome by BOWTIE2 (Langmead and Salzberg, 2012), percentage is computed respect to non tRNA/rRNA pairs. Mapping rate is computed by the sum of percentage of HISAT mapper pairs and BOWTIE2 mapped pairs. Uniquely mapped indicates number of read pairs that are aligned to the human genome at a single location without ambiguity by HISAT2 (with alignment tag NH:1) or BOWTIE2 (MAPQ = 255). Unique map rate is computed respective to the sum of HISAT mapped pairs and BOWTIE2 mapped pairs. Splice read rate is computed by dividing the number of read pairs with either read being split by HISAT2 aligner (with a N cigar string) to the sum of HISAT mapped pairs and BOWTIE2 mapped pairs.

**Table S15**: Mapping statistics for RNA-seq.

### Supplementary videos

**Video S1**: SON/SRRM2 distribution in the nucleus. Visualization of SON and SRRM2 signal in a HepG2 hepatocellular carcinoma cell by immunofluorescent staining. Blue: Hoechst staining of DNA; green: SRRM2; red: SON, yellow: colocalization. Sequence in video: SON only – SRRM2 only – SON and SRRM2 – SON and SRRM2, colocalized areas highlighted in yellow.

**Video S2**: ArgRS/SRRM2 colocalization in the nucleus. Visualization of ArgRS and SRRM2 nuclear signal in a HepG2 hepatocellular carcinoma cell by immunofluorescent staining. The cell nucleus was identified by Hoechst staining. Blue: Hoechst staining of DNA; red: ArgRS; green: SRRM2; yellow: colocalization between ArgRS and SRRM2. Sequence in video: ArgRS only - SRRM2 only - ArgRS and SRRM2 – ArgRS and SRRM2, colocalized areas highlighted in yellow.

**Video S3**: MetRS/SRRM2 colocalization in the nucleus. Visualization of MetRS and SRRM2 nuclear signal in a HepG2 hepatocellular carcinoma cell by immunofluorescent staining. The cell nucleus was identified by Hoechst staining. Blue: Hoechst staining of DNA; red: MetRS; green: SRRM2; yellow: colocalization between MetRS and SRRM2. Sequence in video: MetRS only - SRRM2 only - MetRS and SRRM2 – MetRS and SRRM2, colocalized areas highlighted in yellow.

**Video S4**: MetRS/ArgRS colocalization in the cell. Visualization of ArgRS and MetRS in a HepG2 hepatocellular carcinoma cell by immunofluorescent staining. The cell nucleus was identified by Hoechst staining. Blue: Hoechst staining of DNA; red: MetRS; green: ArgRS; yellow: colocalization between ArgRS and SRRM2. Sequence in video: MetRS only - ArgRS only - ArgRS and MetRS – ArgRS and MetRS, colocalized areas highlighted in yellow.

**Video S5**: Time-lapse of fluorescence recovery after photobleaching (FRAP) of SRRM2-mVenus fluorescence in 293T cells expressing a non-targeting shRNA (ctrl). Bleaching occurs between frame 4 and 5 (1 s into the video).

**Video S6**: Time-lapse of fluorescence recovery after photobleaching (FRAP) of SRRM2-mVenus fluorescence in 293T cells expressing a shRNA against the 3'UTR of ArgRS (shRARS_1). Bleaching occurs between frame 4 and 5 (1 s into the video).

**Video S7**: Time-lapse of fluorescence recovery after photobleaching (FRAP) of SRRM2-mVenus fluorescence in 293T cells expressing a shRNA against the coding region of ArgRS (shRARS_2). Bleaching occurs between frame 4 and 5 (1 s into the video).
