## Supplementary material for "Arg-tRNA synthetase links inflammatory metabolism to RNA splicing and nuclear trafficking via SRRM2": Resource Table

**RESOURCES TABLE**

| REAGENT | SOURCE | IDENTIFIER |
|---|---|---|
| Antibodies | | |
| ArgRS | Biorbyt | orb247357 |
| ArgRS | Abclonal | A6307 |
| ArgRS | PMID 32817337 | RARS-5 |
| ArgRS | Proteintech | 66815-1-Ig |
| SRRM2 | SCBT | sc-390315 |
| SRRM2 | Life Technologies | PA566827 |
| MetRS | Abcam | ab31541 |
| MetRS | Proteintech | 14829-1-AP |
| GFP | Proteintech | 10087-514 |
| SON | Sigma Aldrich | HPA023535 |
| Coilin | CST | 14168T |
| SFPQ | Proteintech | 15585-1-AP |
| PTBP1 | Proteintech | 12582-1-AP |
| GAPDH | CST | 2118L |
| Histone H3 | CST | 9715S |
| Tubulin | CST | 2144S |
| Anti-Rabbit HRP | Invitrogen | 31464 |
| Anti-Mouse HRP | Invitrogen | PI31432 |
| Anti-Rabbit Alexa 568 | Abcam | ab175696-500ug |
| Anti-Mouse Alexa 488 | Abcam | ab150113-500ug |
| Bacterial and Virus Strains | | |
| DH5alpha | NEB | C2987H |
| Top10 | Invitrogen | C404010 |
| ClearColi | Lucigen | 89428-536 |
| Biological Samples | | |
| Chemicals, Peptides, and Recombinant Proteins | | |
| Marfey's Reagent | Pierce | PI48895 |
| Murine Arginase-1 | This paper | N/A |
| Hoechst 33342 | Adipogen | 50596053 |
| Carbon tetrachloride ($CCl_4$) | Sigma-Aldrich | 270652-100ML |
| 13C6-arginine | Pierce | PI88210 |
| Alamar Blue | Biorad | BUF012A |
| HPG-ALD Polymers | Flintbox UBC | N/A |
| T4 Polynuclotide Kinase, Cloned | Lucigen | P0503K |
| TGIRT-III | Ingex | TGIRT™-III Enzyme |
| Thermostable 5' App DNA/RNA Ligase | New England Biolabs | M0319 |
| Phusion High Fidelity PCR Master Mix with HF Buffer | Thermo Scientific | F531 |
| TGIRT III | Ingex | TGIRT™-III Enzyme |
| Critical Commercial Assays | | |
| Co-IP kit | Pierce | PI26149 |
| mirVana RNA isolation | Life Technologies | AM1560 |
| OneStep RT PCR | Qiagen | 210210 |
| NEBNext Magnesium RNA Fragmentation Module | New England Biolabs | E6150S |
| 5' DNA Adenylation Kit | New England Biolabs | E2610 |

| | | |
|---|---|---|
| NEBNext Ultra II Directional RNA Kit for Illumina | New England Biolabs | E7760 |
| Poly(A)Purist MAG Kit | Invitrogen | AM1922 |
| RNA Clean & Concentrator-5 Kit | Zymo Research | R1013 |
| Oligo Clean & Concentrator Kit | Zymo Research | D4060 |
| MinElute Reaction Cleanup Kit | Qiagen | 28206 |
| Pierce High pH Reversed-Phase Peptide Fractionation kit | Thermo Scientific | 84868 |
| Deposited Data | | |
| TGIRT-seq data | This paper | SRA PRJNA561913 |
| Poly(A) RNA-seq | This paper | GEO GSE165513 |
| ArgRS interactome | This paper | Pride PXD015692 |
| MetRS interactome | Cui et al., 2020 | Pride PXD021527 |
| N-terminomics | This paper | Pride PXD024091 |
| Experimental Models: Cell Lines | | |
| HepG2 | ATCC | HB-8065, verified by ATCC |
| HEK 293T | ATCC | CRL-3216, verified by ATCC |
| Murine embryonic fibroblasts | Dr. Yao Tong Scripps Research | N/A |
| Experimental Models: Organisms/Strains | | |
| C57Bl/6J | Scripps Breeding Colony | N/A |
| Oligonucleotides | | |
| gRNA SRRM2 CCAUGAGACACCGCUCCUCC | Eton Bioscience | N/A |
| shRNA RARS (3'UTR) gtggacacaagcataagtaaa | Eton Bioscience | N/A |
| shRNA RARS (CDS) ggagcagttacaagaagaaaa | Eton Bioscience | N/A |
| All other oligos | Eton Bioscience | N/A |
| TGIRT-seq: starting molecule Read2 oligonucleotides | IDT Bioscience, Xu et al. 2019 | NTT R2R DNA/R2 RNA |
| TGIRT-seq: 5' phosphorylated Read1 oligonucleotide | IDT Bioscience, Xu et al. 2019 | R1R DNA |
| Recombinant DNA | | |
| pSpCas9(BB)-2A-GFP_SRRM2 | This paper | N/A |
| mVenus donor for SRRM2 | This paper | N/A |
| pLKO.1 shRARS_3UTR | This paper | N/A |
| pLKO.1 shRARS_2 | This paper | N/A |
| pSpCas9(BB)-2A-GFP (PX458) | Ran et al., 2013 | Addgene 48138 |
| pLKO.1 shSCR | Saharia et al., 2008 | Addgene 17920 |
| pLKO.1 shSRRM2_1, _2 | Sigma Aldrich | TRCN0000314562, TRCN0000314504 |
| pLKO.1 shMARS_1, _2 | Sigma Aldrich | TRCN0000298212, TRCN0000293830 |
| Software and Algorithms | | |
| Imaris | Bitplane | https://imaris.oxinst.com/ |
| Fiji | Schindelin et al., 2012 | https://imagej.net/Fiji |
| Hisat2 | Kim et al., 2015 | http://daehwankimlab.github.io/hisat2/ |

| | | |
|---|---|---|
| Bowtie2 | (Langmead and Salzberg, 2012 | http://bowtie-bio.sourceforge.net/bowtie2/index.shtml |
| Trimgalore | Felix Krueger | https://www.bioinformatics.babraham.ac.uk/projects/trim_galore/ |
| Subread | Liao et al., 2014 | https://bioconductor.org/packages/release/bioc/html/Rsubread.html |
| DESeq2 | Love et al., 2014 | http://bioconductor.org/packages/release/bioc/html/DESeq2.html |
| DEXseq | Anders et al., 2012 | https://bioconductor.org/packages/release/bioc/html/DEXSeq.html |
| Vast-Tools | Irimia et al., 2014 | https://github.com/vastgroup/vast-tools |
| Vast Tools Diff module | Han et al., 2017 | https://github.com/vastgroup/vast-tools |
| VastDB | Tapial et al., 2017 | http://vastdb.crg.eu/wiki/Main_Page |
| Panther | Mi et al., 2019 | http://pantherdb.org/ |
| ShinyGO | Ge et al., 2020) | http://bioinformatics.sdstate.edu/go/ |
| Maxquant | Cox,J. and Mann,M., 2008 | https://www.maxquant.org/ |
| Perseus | Tyanova et al., 2016 | https://maxquant.net/perseus/ |
| GraphPad Prism 7-9 | GraphPad Software LLC | https://www.graphpad.com/ |
| R Studio | R Studio | https://rstudio.com/ |
| Tidyverse | Tidyverse/RStudio | https://www.tidyverse.org/ |
| biomaRt | Durinck S et al., 2009 | https://bioconductor.org/packages/release/bioc/html/biomaRt.html |
| EnhancedVolcano | Blighe K, Rana S, Lewis M (2020). | https://bioconductor.org/packages/release/bioc/html/EnhancedVolcano.html |
| TGIRTseq pipeline | Douglas C. Wu | https://github.com/wckdouglas/tgirt_map |
| ggpubr | Alboukadel Kassambara | https://CRAN.R-project.org/package=ggpubr |
| Subread to DEXSeq | Vivek Bhardwaj | https://github.com/vivekbhr/Subread_to_DEXSeq |
| Other | | |
